# Structures of dynamic interactors at native proteasomes by PhIX-MS and cryoelectron microscopy

**DOI:** 10.1101/2025.07.31.667872

**Authors:** Kitaik Lee, Hitendra Negi, Xiang Chen, Katerina Atallah-Yunes, Sunny Truslow, Rithik E. Castelino, Mary R. Guest, Anthony M. Ciancone, Sergey G. Tarasov, Raj Chari, Kylie J. Walters, Francis J. O’Reilly

## Abstract

Proteasome function depends on a network of transient interactions that remain structurally and functionally unresolved. We developed PhIX-MS (Photo-induced In situ Crosslinking-Mass Spectrometry), a structural proteomics workflow that stabilizes transient interactions in cells by UV-activated crosslinking to capture topological information. Applying PhIX-MS with cryo-electron microscopy (cryo-EM), we mapped redox sensor TXNL1 at the proteasome regulatory particle (RP), placing its PITH domain above deubiquitinase RPN11 and resolving its dynamic thioredoxin domain near RPN2/PSMD1 and RPN13/ADRM1, ideally located to reduce substrates prior to proteolysis. We also resolved chaperone PSMD5 bound to RP without the proteolytic core particle (CP) where its C-terminus inserts into the ATPase pore blocking CP binding. PhIX-MS and AlphaFold modeling tether ubiquitin ligase UBE3C/Hul5 along the RP placing its catalytic site above the RPN11 active site, enabling their coupled activities. Our integrative approach enables the localization of native, low-affinity protein interactions and is broadly applicable to dynamic macromolecular assemblies.

## Introduction

The ubiquitin-proteasome system (UPS) is the primary mechanism for targeted protein degradation, removing damaged, misfolded, and/or regulatory proteins that require tightly controlled lifespans^1^. Proteolysis occurs at a hollow, cylindrical 20S core particle (CP) composed of four heptameric rings, with two interior β-rings (PSMB1-7) and two outer α-rings (PSMA1-7). Degradation by proteasomes is highly regulated and coordinated with ∼1000 enzymes involved in post-translational modification of substrates with ubiquitin^2,3^. The proteasome and its associated ubiquitination machinery are established therapeutic targets, with CP inhibitors standard of care for hematological cancers^4^.

To degrade ubiquitinated proteins, the CP is capped at one end (26S) or both ends (30S) by a 19S regulatory particle (RP)^5,6^, which orchestrates substrate recognition, deubiquitination, unfolding, and translocation into the CP and can be biochemically separated into base and lid subcomplexes^7,8^. The base includes ubiquitin receptors (RPN1, RPN10, and RPN13) to bind ubiquitinated substrates^7^ and a dynamic heterohexameric AAA+ ATPase ring (RPT1-6) that unfolds and translocates substrates into the CP^1^. The lid proteins fan out along the base and includes the essential deubiquitinase RPN11/PSMD14^9^. The proteasome binds transiently to many proteins that are not dedicated subunits, including ubiquitination machinery^10–14^, such as the ubiquitin HECT E3 ligase paralogs UBE3A/E6AP and UBE3C/HUL5.

30S/26S proteasomes can specialize across tissue and cell types by incorporating proteins only expressed in a specific tissue^15^. These specialized binders allow proteasome activity to adapt to cell type or context-specific needs. Alternative proteasome regulators can replace the RP to confer specialized functions in response to cellular stress or immune signaling^16,17^. Notably, bacterial infection was found to trigger changes in proteasome composition and function by recruiting PSME3 to the CP to generate cleaved peptides that enhance antimicrobial activity^18^. However, these interactions are notoriously dynamic, resulting in technical challenges for capturing the architecture of these varied proteasome complexes, particularly in their physiological context.

Mass spectrometry has been foundational to defining the dynamic proteasome interactome, including by characterizing proteasomes isolated from cells with a purification handle^13^, taking advantage of cross-linking approaches^19–21^, affinity pulldown^22,23^, differential ultracentrifugation^24^, and proximity labeling^25^. Many of these approaches lack topological information about the protein interactions that are discovered, so cannot readily direct further structure/function experiments. We therefore developed PhIX-MS (Photo-induced In situ Crosslinking Mass Spectrometry), a structural proteomics strategy to characterize the endogenous proteasome interactome directly in cells. Photo-activated crosslinkers, when activated by UV-light are reactive for only nanoseconds and therefore limit crosslinking to true interfaces^26–28^. We combine in situ photo-crosslinking to covalently stabilize transient interactions, and after affinity enrichment, we use crosslinking mass spectrometry to identify the crosslinked residues. These crosslinks serve as molecular rulers that guide targeted structural analyses by AI-assisted structure modelling to provide an atlas of the dynamic proteasome interactome. These approaches revealed the binding sites and structural context for several regulators in the colon cancer HCT116 cell line. We focus on UBE3C/HUL5, which plays a critical role in preventing substrate escape from proteasomes prior to their degradation^29–31^, revealing its location around the substrate entry channel and above RPN11. We complement PhIX-MS with cryo-electron microscopy (cryo-EM) to resolve oxidative stress factor TXNL1, with its thioredoxin domain at RPN2, able to reduce substrates prior to their translocation and proteolysis. PhIX-MS with cryo-EM also resolves proteasome chaperone PSMD5/S5b, revealing how it promotes RP assembly and restricts 26S/30S proteasome formation. In addition to resolving how these three factors bind and act at proteasomes, we provide a structural blueprint of the proteasome interactome in human cells and establish a broadly applicable platform to study protein complexes in their native environment.

## Results

### PhIX-MS – a strategy to trap the interactome of proteasome complexes

To isolate proteasomes for mass spectrometry analyses, we used HCT116 cells edited to include a biotinylated tag-TEV purification handle fused to endogenously expressed proteins at different positions within the proteasome complex, including the RP lid, RP base, or CP. We had previously engineered a cell line with the handle in RPN1^32^, a ubiquitin-binding^33,34^ subunit of the RP base subcomplex that is positioned against the ATPase ring (Figure 1A). Here, we generated two additional cell lines (Extended Data 1) with the biotin-TEV handle fused to a lid component (RPN11, an essential deubiquitinase^9^ located near the substrate entrance channel of the ATPase ring^35^) and to CP subunit PSMB4/β4 (Figure 1A and S1A). Proteasomes from RPN1-tagged, RPN11-tagged and PSMB4-tagged cells were readily purified by streptavidin beads with the expected SDS-PAGE subunit profile (Figure S1B). The purified proteasome complexes were proteolytically active (Figure S1C), with RPN11 tagging indicating equivalent activity for 26S and 30S complexes, whereas RPN1 or PSMB4 tagging yielded predominately active 30S or CP, respectively.

**Figure 1.**
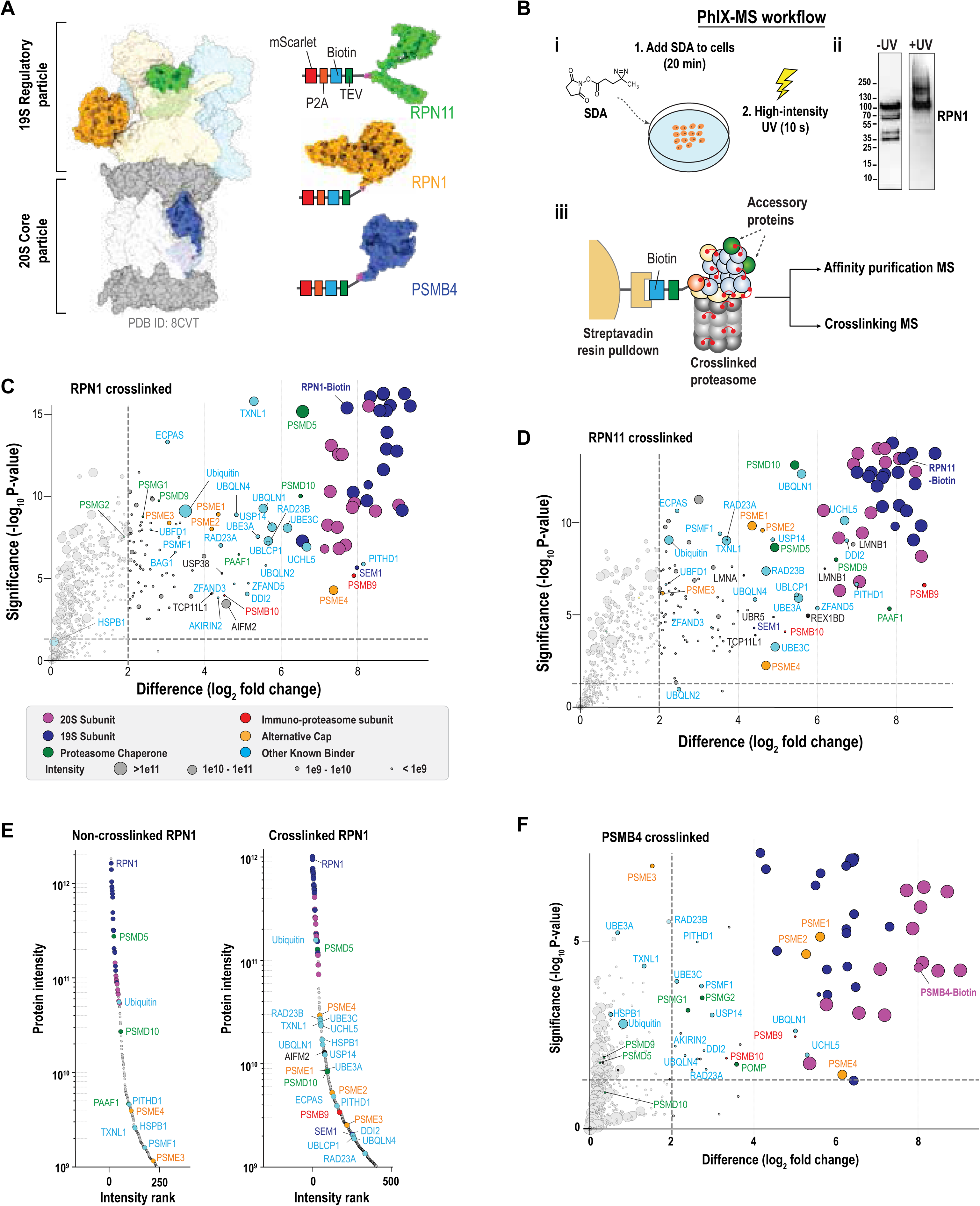
PhIX-MS approach. (**A)** Surface rendering of the 26S proteasome to highlight subunits engineered to include a biotinylated tag. (**B)** PhIX-MS workflow to capture structural information on the proteasome interactome in situ. i) Protein complexes are trapped inside cells by addition of crosslinking agent SDA. Following 20 minutes of incubation, high intensity UV-irradiation is applied for 10 seconds. ii) Immunoprobing lysates for RPN1 indicates higher molecular weight protein complexes following crosslinking of cells. iii) Crosslinked proteasomes are pulled down on streptavidin beads. On-bead digestion is used to identify enriched proteins by AP-MS. Proteasomes are eluted with TEV for analysis by crosslinking MS. The detailed workflow and the crosslinking validation is shown in Figure S1D. **(C)** AP-MS plots of pulldown of biotin-tagged RPN1 from crosslinked cells. Crosslinked wild-type cells are used as the control. Proteins are color-coded by category—core particle (CP, pink), regulatory particle (RP, dark blue), immunoproteasome subunit (red), alternative cap component (orange), proteasome chaperone (green), or other established proteasome binders (cyan). The horizontal dashed line denotes statistical significance (BH-adjusted P < 0.05), and the vertical dashed lines mark log2-fold changes in abundance between tagged and wild-type pulldowns. Dot size represents label-free quantification (LFQ) intensity for each protein. **(D)** Same as for ‘C’ but with of biotin-tagged RPN11. **(E)** Rank intensity plots showing protein abundance (Intensity from MSFragger search) of protein abundances from biotin-tagged RPN1 pulldowns, both from crosslinked and non-crosslinked cells. **(F)** Same as for ‘C’ but with of biotin-tagged PSMB4.

We next developed a strategy, which we name PhIX-MS (Photo-induced In situ Crosslinking Mass Spectrometry, Figure 1B and S1D), to trap transiently or weakly bound proteasome interactors. Prior to lysis, we treat cells with cell-permeable, heterobifunctional crosslinker succinimidyl 4,4’-azipentanoate (SDA), which reacts predominately with primary amines (including lysine and N-termini of proteins), and then photo-activate it to form short-lived (nanoseconds) intermediates that react with any nearby (Cα to Cα, < 27Å) residue^26,27^. This crosslinking mechanism enables time-resolved and spatially constrained mapping of protein-protein interactions in situ. Following purification by the biotin handle, label-free quantitative mass spectrometry (affinity purified-mass spectrometry, AP-MS) was used to identify crosslinked proteins.

By comparing pulldowns without or with crosslinking (Figure S2A and S2B for RPN1, Figure S2C and S2D for RPN11), we found this approach to significantly enriched known proteasome interactors (Figure 1C-1D, Extended Data 2). The crosslinking kept fidelity to native proteasome interactions, as shown by the enrichment of very few typical AP-MS contaminants and abundant cellular proteins^36^. Even with the very mild crosslinking performed here, the abundance of the proteasome-interacting proteins in these RP pulldowns increased dramatically compared to non-crosslinked pulldowns (Figure 1E, S2E, and S2F). Ubiquitin is particularly increased following crosslinking with the purification handle on RPN1, perhaps because it itself is a ubiquitin-binding substrate receptor^33^, and correspondingly, so too are proteins that interact with proteasomes in a ubiquitin-dependent manner, including ubiquitin shuttle factors^37^ RAD23B, RAD23A, UBQLN1, UBQLN4, DDI2, deubiquitinase USP14^38^ and ubiquitin E3 ligase UBE3C^38^ (Figure 1E).

UBE3C is not detected in non-crosslinked pulldowns but represents 4.6% and 3.0% of the RP label-free quantification (LFQ) intensity in the RPN11 and RPN1 pulldowns, respectively (Figure S2E and S2F). Strikingly, alternative regulatory caps including PSME4 (PA200) and PSME1/2/3 (PA28α/β/γ) were strongly enriched in crosslinked samples (Figure S2E, S2F). PSME4 was present at 30% (RPN1, Figure S2G, top) or 17% (RPN11) of the CP LFQ intensity identified in the crosslinked pulldowns (Figure S2G, top). This finding supports the presence of a large percentage of hybrid proteasome complexes, such as 19S-20S-PSME4 and 19S-20S-PA28, in cells at an abundance larger than previously represented, most likely due to earlier studies lacking in situ stabilization^23^. Given that our crosslinking conditions were intentionally mild, even these values may underestimate the true abundance of hybrid assemblies.

The PSMB4 pulldowns enriched the CP chaperones PSMG1, PSMG2, and POMP, which were not pulled down by tagged RPN11 or RPN1 (Figure 1F and S2H). Conversely, the RP chaperones PSMD9/p27, PSMD5/S5B, PSMD10/p28, and PAAF are not enriched in the CP pulldowns (Figure 1F, S2I, and S2J). Alternate 20S CP caps PSME1, PSME2, and PSME4 were strongly enriched in the PSMB4 pulldowns, which did not suggest any further alternate caps beyond those previously reported (Figure 1F, S2G, bottom).

Altogether, this photo-crosslinking approach traps transient interactors and chaperones at proteasome complexes with minimal non-specific background and reveals a greater than expected abundance of hybrid proteasomes.

### PhIX-MS-derived distance restraints filter AlphaFold predictions to identify novel interfaces

We identified crosslinked residue pairs in these pulldowns by crosslinking mass spectrometry. The RPN1- and RPN11-pulldown datasets combined yielded 2236 inter-protein links describing 276 protein-protein interactions (PPIs) (Extended Data 3). Looking at crosslinks only between canonical proteasome subunits, we find 562 crosslinks from the RPN1-purified and 1,697 crosslinks from RPN11-purified complexes (Figure S3A). The majority of these intra-proteasome crosslinks are localized within the RP (Extended Data 3), as expected from the location of the affinity tags (Figure 1A). The RP undergoes conformational switching that is dependent on the presence and status of ubiquitinated substrates, with large changes previously observed for RPN1 and RPN11, as previously summarized^1^. Mapping the crosslinking-based distance information onto any one of the available structures yields clusters of crosslinked residues that are overlength (>27 Å Cα-Cα), suggesting alternative conformational states (Figure S3B). However, 93.5% of the crosslink distances were satisfied on at least one of the 20 available proteasome structures (Figure S3C, Extended Data 4, PDB IDs: 6MSB, 6MSD, 6MSE, 6MSG, 6MSH, 6MSJ, 6MSK, 7W37, 7W38, 7W39, 7W3A, 7W3B, 7W3C, 7W3F, 7W3G, 7W3H, 7W3I, 7W3J, 7W3K, and 7W3M). Different clusters of crosslinks representing different conformational states is apparent by inspection of the position of RPN1 relative to the ATPase ring in S_B_^USP14^ (PDB 7W3I) and E_A1_^UBL^ (PDB 7W37) (Figure S3D). We therefore concluded that the purified proteasome samples previously studied closely recapitulate the native in-cell conformational ensembles, but also that additional conformations may exist. One area where the crosslinking data suggest a previously unobserved configuration is between RPN8 and RPN9 (Figure S3E).

Despite the large number of intra-proteasome crosslinks detected, only a limited number of crosslinks were observed between canonical proteasome subunits and transient interactors (Figure S3F). Correspondingly, the crosslinking MS dataset from the PSMB4 pulldown identified very few RP binders (Figure S3G). This sparsity may reflect low abundance, structural flexibility, and the relatively mild crosslinking conditions used. The 276 PPIs in the combined RPN1 and RPN11 datasets have an estimate FDR of 7.7% based on our target-decoy approach (Extended Data 3). These false PPI matches are unlikely to be between intrinsic proteasome subunits, and we therefore assumed them to concentrate among the potential proteasome interactors. To remove likely false matches, we filtered out the crosslinking interactions for proteins that were enriched in the AP-MS samples (Figure 1C, 1D, and 1F). This filtering step reduced the number of proteins identified with direct crosslinks to intrinsic proteasome subunits from 30 (Figure S3F) to 16 (Figure 2A). Of these, experimental structures are available at proteasomes for PSME1/2^39^ (Figure S4A, PDB 7DR6), ubiquitin chains (Figure S4B, PDB 8JTI), and deubiquitinase USP14^35,40^ (Figure S4C, PDB 7W37), all of which are consistent with the measured crosslinks. Similarly, the structure of UCHL5^41,42^ with the DEUBAD domain from RPN13, to which it binds, and ubiquitin is consistent with our crosslinking data (Figure S4D, PDB 4UEL). These findings provide confidence that the determined structures are adopted in the cellular context and that our crosslinking data can be used to provide high-fidelity structural information.

**Figure 2.**
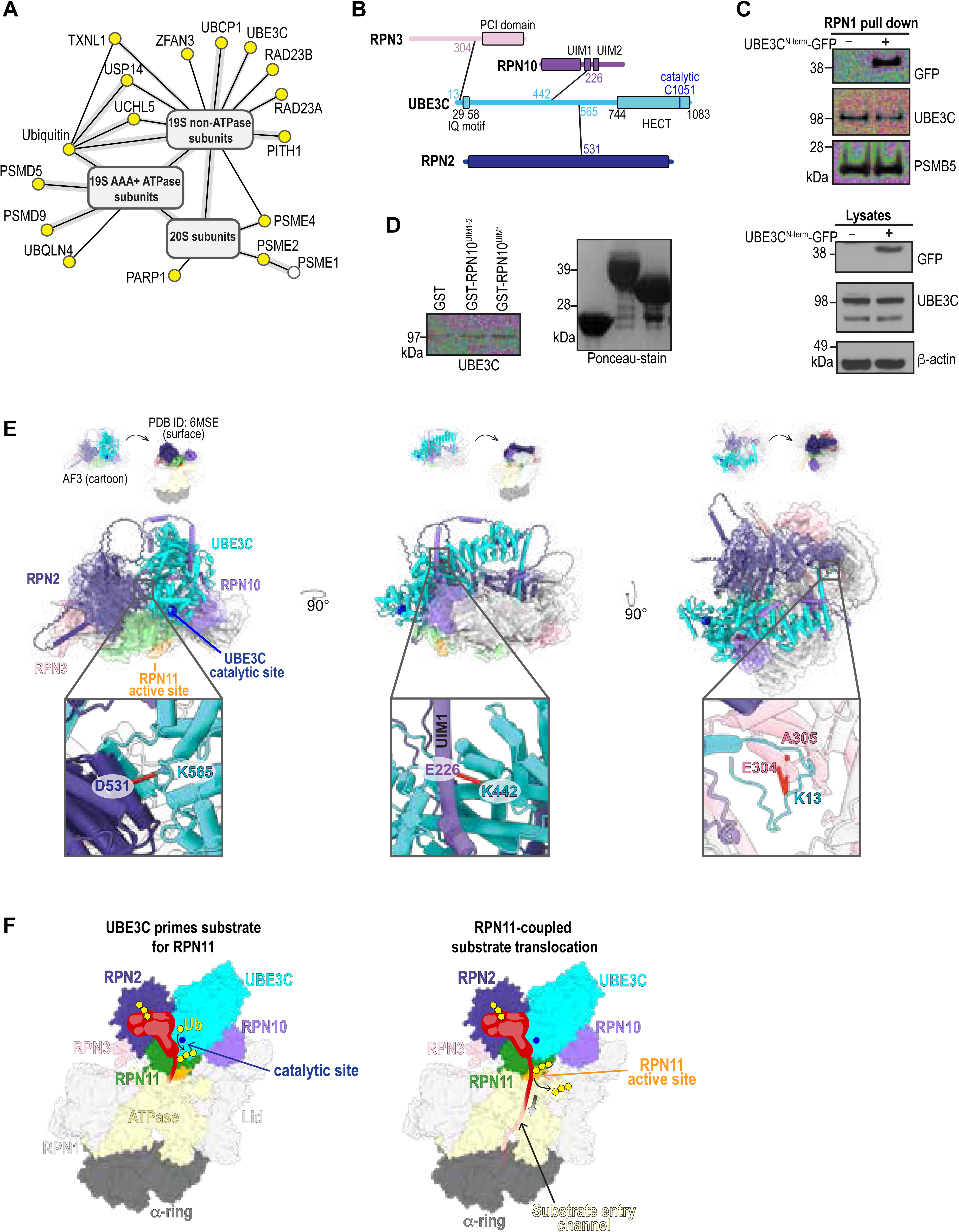
PhIX-MS derived crosslinks and structural modelling localize UBE3C on the RP. **(A**) Crosslink-based protein-protein interaction network of proteins bound to the proteasome filtered for those enriched by AP-MS. **(B)** Protein sequence location of crosslinked residue pairs between UBE3C and proteasome subunits with domains annotated. The UBE3C catalytic cysteine (C1051) is also labeled. (**C)** Immunoblots for UBE3C and GFP as indicated of RPN1-tagged cell lysates (bottom) or following pulldown with streptavidin resin (top) without (−) or with (+) overexpression of UBE3C^N-term^-GFP. PSMB5 and β-actin were used as controls. **(D)** Immunoblot for UBE3C of samples from a pulldown experiment in which glutathione sepharose resin bound to GST-RPN10^UIM1-2^, GST-RPN10^UIM1^, or GST (control) was incubated with HCT116 whole cell lysates, washed to remove unbound proteins, and the proteins then eluted by 2X SDS loading dye prior to SDS-PAGE and immunoprobing with anti-UBE3C antibodies (left). As a loading control for the bound proteins, the PVDF membrane was stained with Ponceau reagent and imaged before immunoprobing (right). **(E)** AlphaFold3 model of UBE3C bound to proteasomal subunits RPN2, RPN3, RPN8, RPN9, RPN10, RPN11, and RPN12 (top, ribbon) docked onto a human 26S proteasome cryo-EM structure in the substrate engaged state (PDB 6MSE, top, surface) showing in the middle panels RPN2 (indigo), RPN3 (pale pink), RPN10 (purple), RPN11 (green) highlighting its active site (orange), and UBE3C (cyan, catalytic cysteine C1051, indigo sphere) as ribbon (AlphaFold3) or surface (PDB 6MSE) displays. Enlarged ribbon diagrams are included below of boxed regions to highlight residue pairs identified by PhIX-MS, including UBE3C K565 to RPN2 D531 (left), UBE3C K442 to RPN10 E226 (middle), and UBE3C K13 to RPN3 E304 and A305 (right). (F) Cartoon of coordinated UBE3C and RPN11 activities for processive translocation of a substrate into the CP. UBE3C (cyan) is shown at the RPN2-RPN10-RPN11 interface adding ubiquitin (yellow) to a proteasome-bound substrate (red, left) priming it for deubiquitination by RPN11 (green with catalytic region in orange) thereby promoting substrate translocation through the ATPase substrate entry channel (right). The model structure is generated from that in panel (E) and follows the same coloring scheme.

We used AlphaFold2-Multimer^43^ to predict the binding sites of the proteins identified as directly crosslinked to the RP. As proteins may have additional interfaces with subunits of the proteasome other than those found crosslinked, we predicted pairwise structural models to the 19 established RP subunits, for a total of 270 potential binary pairs (Extended Data 5). For each protein-subunit pair, five models were generated, and interface confidence was evaluated using model-confidence scores averaged across all five models^44^. Applying a stringent cutoff of 0.65 for average model-confidence score gives six novel interfaces for UBLCP1-RPN1, PSMD5-RPT1, UBE3C-RPN2, RAD23A-RPN1, RAD23B-RPN1, PITHD1-RPN10 (Extended Data 5, Figure S4E). AlphaFold predicts the phosphatase UBLCP1 to use its ubiquitin-like (UBL) domain to bind to the RPN1 T2 site (Figure S4F) where USP14 UBL binds^33^. This binding interface is supported by eight unique crosslinks (Figure S4F). RAD23A (Figure S4G) and RAD23B (Figure S4H) were predicted by AlphaFold to bind to the T1 site, as previously shown in *S. cerevisiae*^33,45,46^, and a crosslink was identified between the RPN1 T1 and T2 sites that matches to but cannot resolve RAD23A or RAD23B UBL (Figure S4E). These in cellulo data suggest that the RPN1 T1 – T2 site may be a high traffic region in the proteasome, contributing to RPN1 dynamics and dictating functionality by competitive occupancy. This phenomenon may in turn drive the timing of fluctuations between the various proteasome configurations and activities.

PITHD1, the RP chaperone PSMD5, and the ubiquitin ligase UBE3C yielded high-confidence AlphaFold models that suggest their binding sites on the proteasome. PITHD1 is predicted to bind to the RPN10 VWA (von Willebrand factor type A) domain (Figure S4I) and along with a crosslink to nearby RPN2, this binding location agrees with a recent preprint that reports the structure of PITHD1 at RPN10 and RPN2 on the proteasome^47^. The PSMD5 predicted structure with ATPase subunit RPT1 had the highest average ipTM (interface predicted Template Modeling) score (0.90) across all predicted interfaces and is supported by four crosslinks (Figure S4J).

In summary, PhIX-MS data and AlphaFold modeling suggest that the RPN1 T1-T2 site is a high traffic exchange region for UBL domains in UBLCP1, RAD23A/B, and USP14 and crosslinks can provide experimental validation to filter AlphaFold predictions.

### UBE3C is positioned along the upper rim of the RP poised above RPN11

UBE3C is a HECT-type ubiquitin E3 ligase with an unknown binding site at proteasomes, where it ubiquitinates protein substrates to promote proteasome processivity, facilitating complete degradation and preventing stalling and/or premature substrate release^29–31^. We find UBE3C crosslinked to three distinct RP subunits, RPN2, RPN3, and RPN10, and that these crosslinks span K13 – K565 (Figure 2B). AlphaFold2 confidently (ipTM score, 0.78) predicts a complexed structure of UBE3C and RPN2 that is supported by the crosslinking data (Figure 2B and S4K), and a lower confidence interface (average ipTM = 0.5) between the disordered N-terminal tail of UBE3C and RPN3, which is also supported by two crosslinks (Figure S4L). This latter finding is consistent with a previous report that the UBE3C N-terminal region is vital for its proteasome association^48^. To experimentally validate the importance of the disordered UBE3C N-terminus, we expressed an N-terminal fragment of UBE3C fused to N-terminus of GFP (UBE3C^N-term^-GFP) in RPN1-tagged cells and tested whether UBE3C^N-term^-GFP can displace endogenous UBE3C from proteasomes. Indeed, endogenous UBE3C was reduced at RPN1-purified proteasome following expression of UBE3C^N-term^-GFP (Figure 2C).

One of the five UBE3C-RPN10 predicted structures suggested UBE3C interaction with the RPN10 UIM1 region (Figure S4M), as indicated by a crosslink between UBE3C K442 and RPN10 E226 that was present in both RPN11 and PSMB4 pulldowns (Figure 2B, S3G, Extended Data 3). To test further whether the RPN10 UIM region is involved in UBE3C binding, we expressed and purified from *E. coli* GST-RPN10^UIM1-2^ (203 – 310), which contains both UIMs, and GST-RPN10^UIM1^ (196 – 272). GST-RPN10^UIM1^ and GST-RPN10^UIM1-2^ each bound modestly to UBE3C in a pull-down assay with GST as a control (Figure 2D and Extended Data 6), offering further support for this interaction.

To complement these findings, we generated a larger model with AlphaFold3 containing UBE3C (cyan), RPN2 (indigo), RPN3 (pink), RPN8 (white), RPN9 (white), RPN10 (purple), RPN11 (green), and RPN12 (white) and selected a structure that recovered all three suggested interaction interfaces (Figure 2E and S4N). Aligning the RP subunits on a proteasome substrate-processing structure (PDB 6MSE) indicated reasonable overlap (Figure S4O and S4P) and fitting of UBE3C without steric clashes (Figure 2E). The model and PhIX-MS data place the N-terminal end of UBE3C at RPN3 from where it extends across RPN2 and RPN10 to position its catalytic cysteine (C1051) above the RPN11 active site (Figure 2E). RPN11 is positioned above the ATPase channel in substrate-processing proteasomes (Figure 1A and 2E), where it removes ubiquitin chains en bloc^49^ with its cleavage activity enhanced as the substrate translocates into the ATPase ring^50^. No crosslinks were observed to residues in the UBE3C HECT domain (Figure 2B), suggesting that this catalytic domain is dynamic at proteasomes. We propose that UBE3C dynamically ubiquitinates substrates, priming them for RPN11 deubiquitination as they translocate into the ATPase channel (Figure 2F). This model provides an explanation for why UBE3C is required for processive proteasomal degradation of substrates.

### Proteasome binding mechanism of TXNL1 by cryo-EM

To complement our MS data with high-resolution structural information, we collected cryo-EM data on RPN1-purified proteasomes (Figure S5A-S5W). We were unable to observe UBE3C at proteasomes in this dataset but readily identified TXNL1 by selecting for particles with additional density near RPN2 where the crosslinking data indicated interaction (Figure 3A and S5A). This effort resolved two distinct structural states with the TXNL1 PITH domain observable at 26S proteasomes, one at 3.82 Å resolution with an opened CP gate (26S^TXNL1-1^ Figure 3B) and the other at 4.0 Å resolution with a closed CP gate (26S^TXNL1-2^, Figure 3C). In both cases, TXNL1 is positioned between RPN2 and the RPN10 VWA domain with its C-terminus extending across RPN11 and ending at the oligonucleotide/oligosaccharide-binding fold (OB) ring of the ATPase (Figure 3B, 3C, and S6A). An opened gate TXNL1-bound state resolving the PITH domain was identified in two other studies^51,52^, and akin to these structures, we observe C-terminal H289 of TXNL1 coordinated to the RPN11 Zn^2+^, which is also coordinated by RPN11 H113, H115, and D126 (Figure 3D, S6A, and S6B). We note that PITHD1 was also reported to bind to this location in a recent pre-print, with a longer C-terminal tail that extends through the ATPase pore^47^.

**Figure 3.**
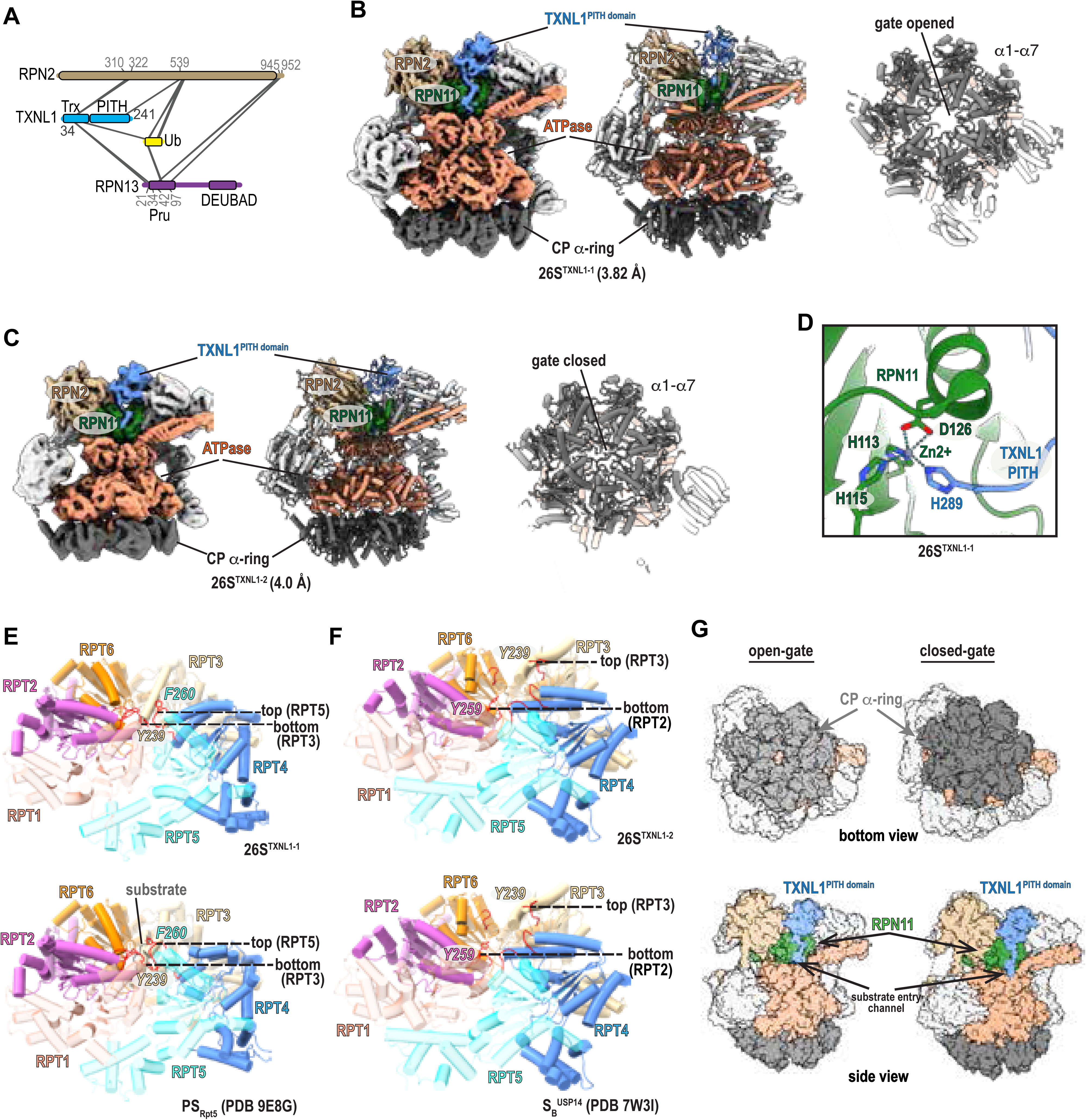
Structures of 26S proteasomes with TXNL1 determined by cryo-EM. **(A**) Protein sequence location of crosslinked residue pairs between TXNL1 and proteasome subunits or ubiquitin. (**B)** Cryo-EM density map (3.82 Å, left) and corresponding ribbon diagram (middle) of 26S^TXNL1-1^. The CP α-ring (dark grey), ATPase (light orange), RPN2 (beige), RPN11(green) and TXNL1 PITH domain (blue) are highlighted and labeled, with the other proteasome subunits colored light grey. The right panel shows a bottom view of the CP α-ring and illustrates the opened gate CP conformation. (**C)** Cryo-EM density map (4.0 Å, left) and corresponding ribbon diagram (middle) of 26S^TXNL1-2^ colored as in (B). The right panel shows the bottom view of the CP α-ring and illustrates the closed gate CP conformation. (**D)** Expanded view of 26S^TXNL1-1^ to display TXNL1 PITH domain and C-terminal residue H289 coordination with the RPN11 Zn^2+^ (purple sphere) and its additional coordination with H113, H115 and D126 of RPN11. (**E and F)** Expanded views of the ATPase large AAA+ subdomains in aligned 26S^TXNL1-1^ (E, top) and PS_Rpt5_ (PDB 9E8G, E, bottom), or 26S^TXNL1-2^ (F, top) and S_B_^USP14^ (PDB 9E8G, F, bottom). RPT subunits are displayed as ribbon diagrams and colored in maroon (RPT1), magenta (RPT2), orange (RPT6), beige (RPT3), light blue (RPT4), and cyan (RPT5), with the pore-1 loops colored in red and the sidechain of the central Tyr/Phe residues in pore-1 loops displayed. The Cα positions of the top and bottom Tyr/Phe residues in pore-1 loops are highlighted and labeled by black dash lines. **(G)** Cartoon depicting the TXNL1 PITH domain (blue) at the proteasome RP with an open (left) or closed (right) CP α-ring (grey). A cutaway view looking through the α-ring to the RP (top) and side view (bottom) highlighting RPN11 (green), the ATPase ring (pink), and the substrate entry channel is displayed. Figure 3B-3G were prepared using UCSF ChimeraX.

Comparison with available PDB files from previous studies indicates the ATPase large AAA+ subdomains in 26S^TXNL1-1^ to adopt a spiral-staircase arrangement similar to the substrate-processing state PS_Rpt5_ (PDB 9E8G). In both cases, RPT5 is at the top of the staircase and RPT3 at the bottom (Figure 3E), with less interleaving of RPT4 with its neighboring subunits (Figure S6C). In contrast to PS_Rpt5_ however, 26S^TXNL1-1^ lacks substrate and in turn, the TXNL1 PITH and RPN10 VWA domains are laterally shifted toward the translocation channel causing the RPT4:RPT5 coiled coil to be rotated outwards (Figure S6D). By contrast, the spiral-staircase configuration of the ATPase larger AAA+ subdomains in 26S^TXNL1-2^ adopts a typical resting-state conformation, with RPT3 at the top, RPT2 at the bottom (Figure 3F), and RPT6 mimicking a connecting seam (Figure S6E). 26S^TXNL1-2^ aligns more closely to USP14-bound 26S proteasomes (Figure 3F) although TXNL1 is not present in this complex^40^ (Figure S6F) and USP14 is not observed in 26S^TXNL1-2^. In summary, we find TXNL1 to bind to 26S proteasomes with both an opened and closed CP α-ring, indicating that it can be at both resting and substrate processing proteasomes (Figure 3G).

### PhIX-MS helps position the TXNL1 thioredoxin domain at RPN2

Dropping from a threshold of 0.04 to 0.012, revealed additional density in 26S^TXNL1-1^ extending from the TXNL1 PITH domain and into which the thioredoxin (Trx) domain fit well (Figure 4A). This placement locates Trx proximal to the RPN2 C-terminus where the RPN13 N-terminal Pru (<u>P</u>leckstrin-like receptor for <u>u</u>biquitin) domain binds^53–55^, consistent with a crosslink between RPN13 K34 and TXNL1 C34 (Figure 3A). RPN13 and the RPN2 C-terminus were not observed, most likely due to the dynamics of this region^55^. We therefore used NMR to test whether RPN13 Pru binds to Trx. Unlabeled TXNL1 Trx domain (TXNL1^Trx^) was added at 5-fold, 10-fold, or 15-fold molar excess to 20 μM ^15^N RPN13 (1-150) preincubated with equimolar quantities of its proteasome binding site (RPN2 (940-953)) and the effects monitored by 2D ^1^H, ^15^N HSQC experiments. Only minor signal shifting was observed, which occurred for amino acids in the helix, such as E118 and H120, and at the N-terminus, including G5 and G14, (Figure 4B). These effect likely reflect non-specific binding, as the Pru helix and N-terminus interacts broadly with peptide sequences^56^. Therefore, we conclude that TXNL1^Trx^ does not form a stable interaction with RPN13 Pru but interacts transiently.

**Figure 4.**
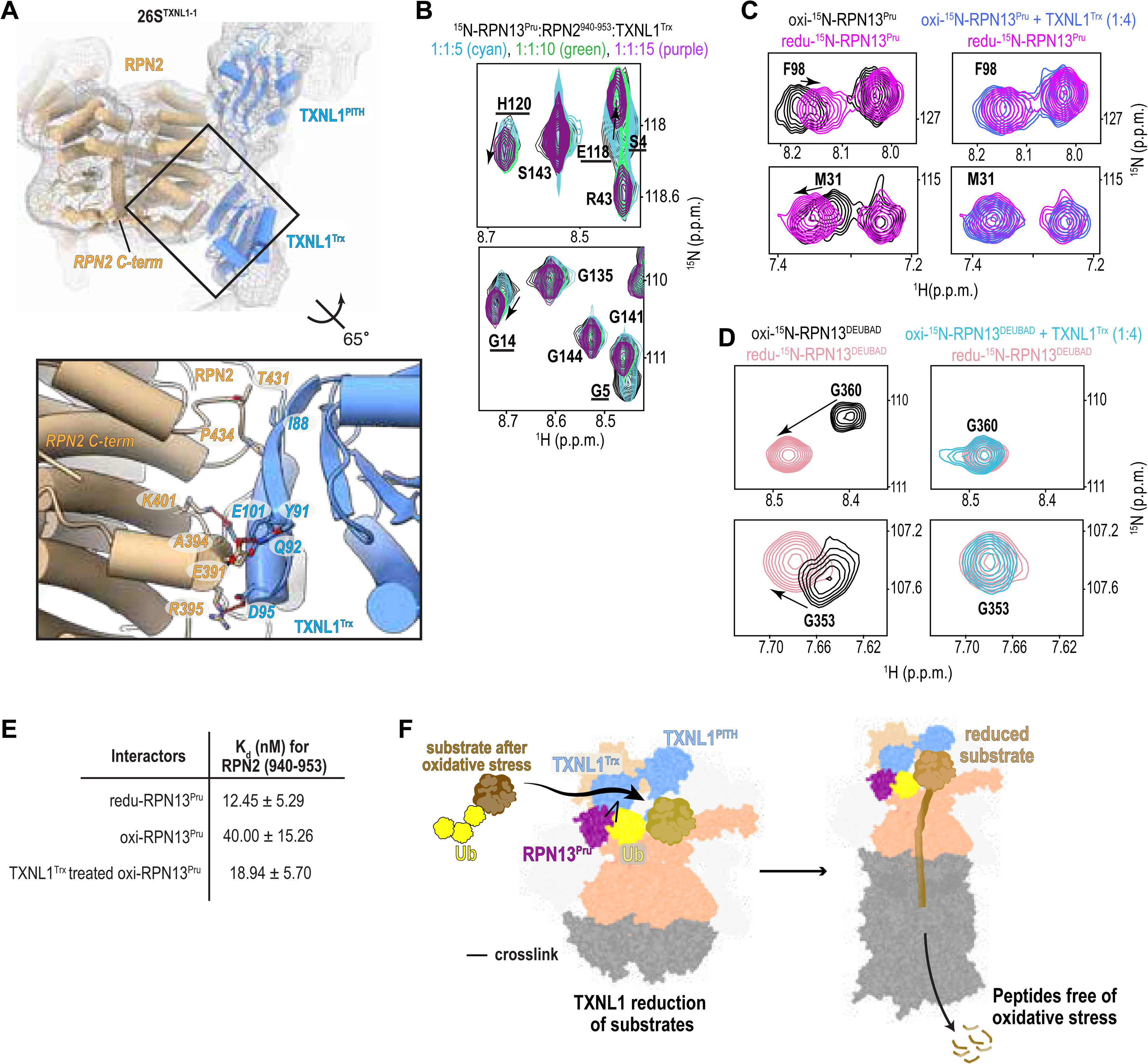
PhIX-MS guides placement of the TXNL1 Trx domain. (**A)** Top panel: Ribbon diagram for an enlarged region of 26S^TXNL1-1^ centered on RPN2 (beige) and TXNL1 (light blue), with the density map displayed in grey. The extreme C-terminus of RPN2 (D953) is not visible and residues E922 – H931 of RPN2 are displayed as spheres. Bottom panel: Expanded structural region to display the contact surface for RPN2 and TXNL1 Trx domain. Sidechains of TXNL1 I88, Y91, Q92, D95, and E101 (light blue) are displayed and labeled, as are RPN2 E391, A394, R395, K401, T431, P434 (beige). Hydrogen bonds are displayed as red lines. Oxygen and nitrogen atoms are colored red and blue, respectively. (**B)** Expanded view of overlaid ^1^H-^15^N HSQC spectra of 20 μM ^15^N-labeled RPN13^Pru^ preincubated with equimolar unlabeled RPN2^940–953^ (black), and with 5-fold (cyan), 10-fold (green), or 15-fold (purple) molar excess unlabeled TXNL1 Trx. RPN13^Pru^ signals that shift or attenuate following addition of unlabeled TXNL1 Trx are labeled and underlined. **(C, D)** Expanded view of overlaid ^1^H-^15^N HSQC spectra of 22 μM ^15^N-labeled RPN13^Pru^ or 25 μM ^15^N-labeled RPN13^DEUBAD^ in reduced (redu, shades of pink) or oxidized (oxi) states without (black) or with 4-fold molar excess TXNL1 Trx domain (shades of blue). The enlarged regions focus on **(C)** residues F98 (top) and M31 (bottom) or **(D)** residues G360 (top) and G353 (bottom). All the spectra were collected in 600 MHz at 10 °C with NMR buffer containing 20 mM sodium phosphate and 50 mM NaCl (pH 6.0); 1 mM DTT was also included in reduced samples. **(E)** Binding affinity, K_d_ (nM), determined by isothermal titration calorimetry for RPN2 peptide (940-953) binding to reduced (redu) or oxidized (oxi) RPN13^Pru^ and oxi-RPN13^Pru^ following incubation with 3-fold molar excess TXNL1^Trx^. TXNL1^Trx^ was removed prior the measurements by size exclusion chromatography. **(F)** Cartoon of the proteasome RP plus CP α-ring (grey) illustrating an incoming substrate (brown) that is oxidized and ubiquitinated (yellow) being reduced by TXNL1 (blue, left) prior to its translocation into the CP (right). The RPN13 Pru domain (purple) is shown bound to the ubiquitin chain (yellow) of the reduced substrate (yellow-brown). Figure 4A and 4F were prepared using UCSF ChimeraX.

Since TXNL1 alleviates oxidative stress by reducing disulfide bonds^57^, we tested whether it is active against oxidized RPN13. The ^15^N-labeled RPN13 Pru or DEUBAD domains were separately oxidized by incubating them on ice for 10 minutes with 0.05 % H_2_O_2_ and following removal of H_2_O_2_ evaluated by MS and 2D NMR to confirm their oxidation. Addition of TXNL1^Trx^ restored the oxidized RPN13 Pru and DEUBAD signals to their reduced position in 2D ^1^H, ^15^N HSQC experiments (Figure 4C-4D and S6G-S6H). We next tested the impact of oxidation and TXNL1^Trx^ restoration on RPN13 Pru affinity for RPN2 (940-953) by using isothermal titration calorimetry (ITC). RPN13 Pru was measured to bind RPN2 with 12.45 ± 5.29 nM affinity, consistent with previous measurements^54^. Following oxidation this affinity remained high but was reduced to 40.00 ± 15.26 nM. We next measured the binding affinity after RPN13 Pru was restored by TXNL1^Trx^, which was removed prior to ITC, to find a value of 18.94 ± 5.70 nM (Figure 4E and S6I). These results indicate that TXNL1^Trx^ can rescue Rpn13 Pru from oxidative stress.

Altogether, we find the TXLN1 PITH domain to cap RPN11 while its Trx domain extends to RPN2 near where RPN13 binds. From this location, TXNL1 is ideally poised to rescue substrates from oxidative stress prior to passage into the substrate entry channel and proteolysis in the CP (Figure 4F). This activity would prevent the production of oxidized peptides.

### PSMD5 binds the ATPase ring to regulate RP maturation

We observed extensive crosslinks between the ATPase ring and PSMD5 in both RPN1 and RPN11 purified proteasomes (Figure 5A), along with a high-confidence AlphaFold model between PSMD5 and RPT1 (Figure S4J). PSMD5 promotes RP assembly^58^ and when upregulated by the NF-κB pathway, inhibits 26S proteasome assembly^59,60^. We performed co-fractionation MS on crosslinked RPN1 pulldowns, which showed that PSMD5 elutes with the intact RP (Figure 5B), consistent with its role in promoting RP assembly and inhibiting 26S formation. Proteomics of these fractions revealed PSMD5 has a similar abundance to all other RP subunits suggesting that most, if not all mature, 19S particles are bound to PSMD5 prior to 26S/30S proteasome formation. To test the hypothesis that PSMD5 levels might regulate 26S proteasome formation, we overexpressed PSMD5 in HCT116 cells prior to RPN1 pulldowns. Cells overexpressing PSMD5 pulled down significantly less 20S CP bound to the 19S RP (Figure 5C). We additionally performed proteasome activity assays in these cell lysates to test the effect of increased PSMD5 levels on proteasome activity. Overexpression of HA-PSMD5 resulted in significant inhibition of 26S proteasome activity (Figure 5D and Extended Data 7), consistent with its proposed role as a negative regulator of 26S proteasome assembly^59^. These findings extend our understanding of PSMD5 as more than a passive chaperone, positioning it as a gatekeeper of RP-CP coupling. Given its stress- and cancer-regulated expression^60^, PSMD5 may act as a rheostat of proteasome capacity in disease contexts, warranting its further investigation as a potential therapeutic target.

**Figure 5.**
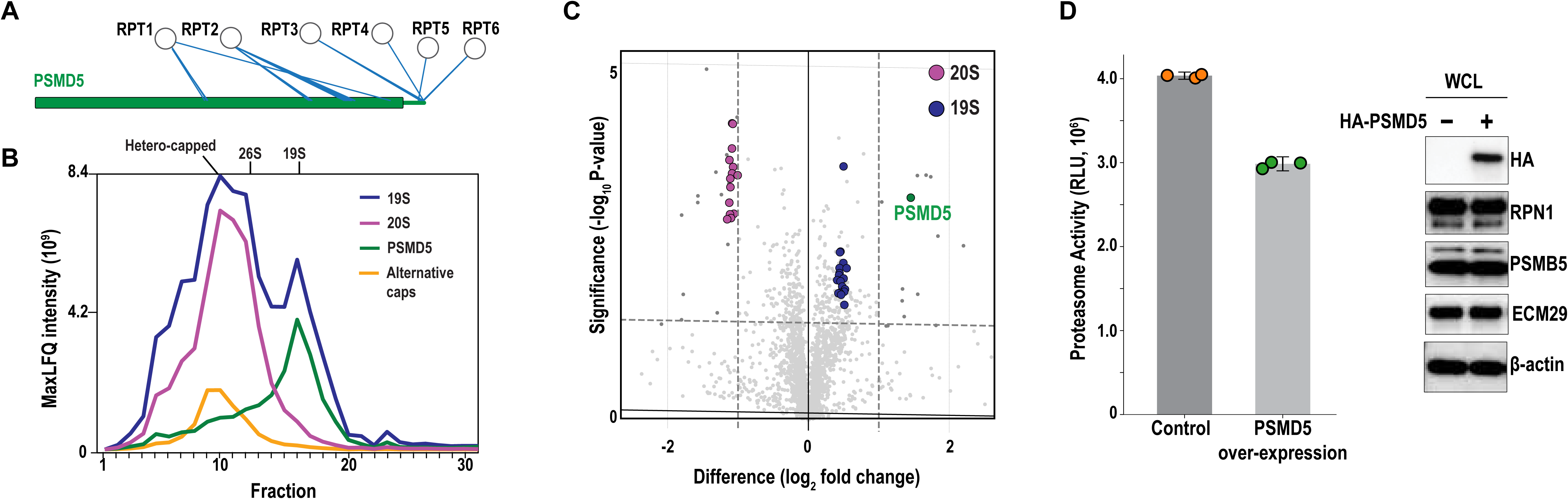
PSMD5 binds to the intact RP. (**A)** PSMD5 sequence plot showing crosslinks to all 6 RP ATPase subunits, including at its disordered C-terminal tail. **(B)** Co-fractionation MS data from an RPN1 pulldown separated by size exclusion chromatography suggests that PSMD5 elutes with the free RP. 20S and alternative cap elution profiles localize 26S and larger proteasome complexes. **(C)** Overexpression of HA-tagged PSMD5 causes reduction of 20S subunits in an RPN1 pulldown. The horizontal dashed line denotes statistical significance (BH-adjusted P < 0.05), and the vertical dashed lines mark log_2_-fold changes in abundance between HA-PSMD5 and control. **(D)** Chymotrypsin-like proteasome activity assays (left) show that HA-PSMD5 overexpression diminishes proteasome activity. Orange and green dots represent individual replicates of the control and HA-PSMD5 overexpression, respectively. Immunoblots (right) confirm HA-PSMD5 expression, with RPN1, PSMB5, ECM29, and β-actin serving as controls.

### Structure of the PSMD5-RP complex by PhIX-MS and cryo-EM

Since the crosslinks suggest PSMD5 interacts with all members of the RP ATPase ring (Figure 5A), we used AlphaFold3 to predict a model of PSMD5 with all six ATPase subunits. We identified a confidently scored model that satisfies 30/31 crosslinks (Figure S7A and S7B). While consistent with prior findings showing direct binding to RPT1 and RPT2^58,60^, the disordered C-terminal tail of PSMD5 is also predicted to insert into the central pore of the ATPase hexamer with a cluster of crosslinks to confirm this prediction (Figure S7C). The main ordered body of PSMD5 is predicted to bind RPT1 and RPT2 in an orientation that would be incompatible with 26S formation (Figure S7B). Indeed, PSMB4 pulldowns of the proteasome CP did not enrich for any PSMD5 protein (Figure S2H-J).

In our cryo-EM dataset, we observed free RP particles (with no CP present) and selected these for further refinement (Figure S5A and S5B). Three distinct states were observed, two with the TXNL1 PITH domain present; one with poor resolution for RPN1, RPT1, and RPT2 (19S^TXNL1-1,ΔRPT1,^ **^Δ^**^RPT2^, Figure 6A left), another in which the TXNL1 C-terminal tail is not visible (19S^TXNL1-2^, Figure 6A middle), and a third without TXNL1 present (19S**^Δ^**^TXNL1-1^, Figure 6A right). 3D classification and 3D variability analyses followed by local refinement masking for 19S^TXNL1-1,ΔRPT1,ΔRPT2^ focused on the RPT1-RPT2-RPT5-RPT6 AAA+ domain and RPN1 revealed density for RPN1 and the AAA+ domain of RPT1 and RPT2 (19S^TXNL1-1^, Figure 6B and S8A). Akin to 26S^TXNL1-1^ and 26S^TXNL1-2^ (Figure S6A), no additional density was observed around the ATPase pore in 19S^TXNL1-1^ or 19S^TXNL1-2^, consistent with the TXNL1 extreme C-terminus culminating at RPN11 (Figure S8B). Extra density did appear however in 19S^TXNL1-1^ where our crosslinking data indicated PSMD5 to bind (Figure 6B). Applying a 10 Å low-pass filter improved the signal-to-noise for this region and we were able to place PSMD5 reliably into this extra density (Figure 6C), satisfying all but one crosslink between RPT1 A289 and PSMD5 K461 (Figure 6D). Applying the same approach on 19S^TXLN1-2^ did not show additional density, suggesting that PSMD5 may be absent at these particles or too dynamic to observe, albeit only 39,459 particles represent this structure (Figure S5B). By contrast, this approach revealed extra density corresponding to PSMD5 at 19S**^Δ^**^TXNL1-2^ (Figure S8C, left with 15 Å low-pass-filtered density on right), which was obtained by 3D variability analyses and masked local refinement of 35,258 particles from 19S^ΔTXNL1-1^ (Figure S5B). Therefore, of the three RP structural states observed (Figure 6B and S8C), two (19S^TXNL1-1^ and 19S**^Δ^**^TXNL1-1^) showed visible PSMD5 following further refinement, one with and one without TXNL1.

**Figure 6.**
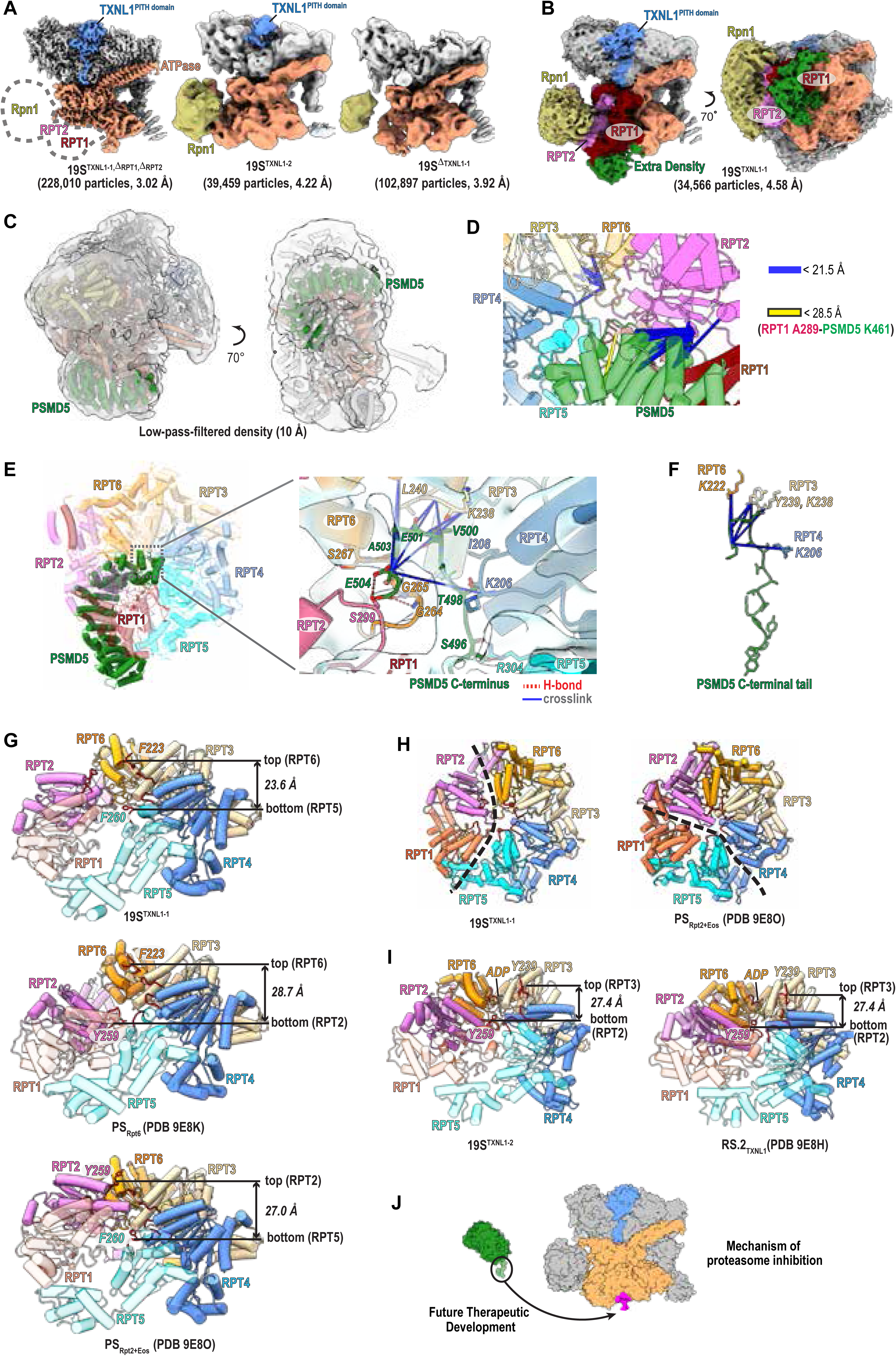
Structures of the 19S RP with TXNL1 and PSMD5 determined by cryo-EM. (**A)** Cryo-EM density map of 19S^TXNL1-1,^ **^Δ^**^RPT1,^ **^Δ^**^RPT2^ (3.02 Å, left), 19S^TXNL1-2^ (4.22 Å, middle), and 19S**^Δ^**^TXNL1-1^ (3.92 Å, right). The ATPase (RPT3, RPT4, RPT5, and RPT6, light orange), RPN1 (beige), and TXNL1(blue) are highlighted and labeled, with the other RP subunits colored light grey. (**B)** Cryo-EM density map of 19S^TXNL1-1^ (4.58 Å, left) and following 70° rotation about the x-axis, using a counterclockwise convention (right). RPT1 (red), RPT2 (magenta), RPN1 (beige), and TXNL1(blue) are highlighted and labeled, with the other RP subunits colored light grey. Extra density adjacent to RPT1/RPT2 is colored in green. (**C)** PSMD5 (green) structure fitted into the extra density adjacent to RPT1 and RPT2 of a low-pass-filtered (10 Å) 19S^TXNL1-1^ map. (**D**) Expanded view of 19S^TXNL1-1^ showing the crosslinks between PSMD5 (green) and RPT subunits as dashed lines and colored blue for distances <21.5 Å and yellow for one between 21.5 and 28.5 Å. (**E**) Left: Ribbon diagram showing the ATPase central channel with the PSMD5 C-terminus inserted. Right: Expanded structural region displaying the contact surface between the PSMD5 C-terminal residues and RPT subunits. Sidechains of PSMD5 S496, T498, V500, E501, A503 and E504 (green) are displayed and labeled, as are RPT2 S299 (magenta), RPT6 G264, G265, S267 (orange), RPT3 K238, L240 (beige), RPT4 K206, I208 (light blue), and RPT5 R304 (cyan). Hydrogen bonds and crosslinked residues are represented by red dashed and blue lines, respectively. **(F)** Expansion of the PSMD5 C-terminal tail with crosslinked residues RPT6 K222, RPT3 Y239 and K238, and RPT4 K206, colored as in (E). **(G and I)** Expanded views of the ATPase large AAA+ subdomains in aligned 19S^TXNL1-1^ (F, top), PS_Rpt6_ (PDB 9E8K, G, middle), and PS_Rpt2+Eos_ (PDB 9E8O, G, bottom), or 19S^TXNL1-2^ (I, left) and RS.2_TXNL1_ (PDB 9E8H, I, right). RPT subunits are displayed as ribbon diagrams and colored in maroon (RPT1), magenta (RPT2), orange (RPT6), beige (RPT3), light blue (RPT4), and cyan (RPT5), with the pore-1 loops colored in red and the sidechain of the central Tyr/Phe residues in pore-1 loops displayed. The Cα positions of the top and bottom Tyr/Phe residues in pore-1 loops are highlighted and labeled by black dashed lines. In (G and I), Cα-Cα distances between the top and bottom Tyr/Phe residues in pore-1 loops are displayed. **(H)** Top views of the ATPase large AAA+ subdomains in aligned 19S^TXNL1-1^ (left) and PS_Rpt2+Eos_ (right), with the same color scheme as used in (G). The reduced interleaving between RPT1/RPT2 and the neighboring RPT subunits in 19S^TXNL1-1^ is highlighted by a black dashed line (left), as is the reduced interleaving between RPT5/RPT1 and the neighboring RPT subunits in PS_Rpt2+Eos_ (right). Figure 6 was prepared using UCSF ChimeraX. **(J)** Model of therapeutic potential to use the ATPase (orange)-interacting PSMD5 (green) C-terminus to develop inhibitors (exemplified in pink) of 26S/30S proteasomes by preventing their assembly.

The structured PSMD5 domain could be fit in 19S^TXNL1-1^ and is positioned against RPT1 while its C-terminus enters the ATPase channel to form extensive interactions involving RPT2, RPT3, RPT4, RPT5, and RPT6 (Figure 6E-F). When aligning this structure with the ATPase-PSMD5 AlphaFold3 model (Figure S8D, grey), poor agreement is displayed for RPT1 (Figure S8D, red), RPT2 (Figure S8D, magenta), RPT5 (Figure S8D, cyan), and RPT6 (Figure S8D, orange). It is possible that the AlphaFold3 model represents an alternative conformational state; however, even if this predicted configuration is not part of the native conformational ensemble, it provided a good starting point for analyses of the experimental data, contributing to the generation of an accurate high-resolution structure.

19S^TXNL1-1^ more closely mimics the substrate-engaged 26S proteasome ATPase state than the resting state, as RPT6 is at the top and RPT5 at the bottom (Figure 6G, top). The arrangement of the hexameric ATPase staircase for 19S^TXNL1-1^ is unique however, with a configuration between previously reported RS_Rpt6_ (PDB 9E8K), where RPT6 is at the top of staircase and RPT2 at the bottom (Figure 6G, middle), and RS_Rpt2+Eos_ (PDB 9E8O), where RPT2 is at the top and RPT5 at the bottom (Figure 6G, bottom). Overall, its ATPase hexamer is flatter compared to RS_Rpt6_ or RS_Rpt2+Eos_, with the distance between the central Phe/Tyr in pore-1 loop decreased from 28.7 Å or 27 Å to 23.6 Å (Figure 6G). This effect is likely driven by PSMD5 inducing reduced interleaving between RPT1/RPT2 and their neighboring subunits (Figure 6H and S8E).

The ATPase configuration of 19S^TXNL1-2^, which lacks observable PSMD5, mimics the resting state observed in 26S proteasomes, with RPT3 at the top, RPT2 at the bottom, and RPT6 at a seam position bound to ADP (Figure 6I). In this structure, the TXNL1 PITH domain is observed but its C-terminal tail not visible (Figure S8F). An analogous resting 26S proteasome state was observed with TXNL1, for which TXNL1 is reported to have lower affinity compared to the substrate-degrading 26S proteasome state^51^. Furthermore, that the ATPase ring of 19S^TXNL1-2^ is not flattened led us to conclude that PSMD5 is not present, rather than not visible due to dynamics. It is possible however that this state was generated during sample handling from breakdown of 26S/30S proteasomes into RP and CP complexes.

Altogether, we find that PSMD5 interacts with RP in a manner that sterically occludes the CP and that engagement of its C-terminal tail into the ATPase ring weakens subunit interleaving to generate a flatter staircase structure.

## Discussion

In this study, we present an integrative structural and proteomic analysis of the proteasome interactome in human cancer cells, combining PhIX-MS, cryo-EM, and computational modeling. By applying this integrative approach to endogenously tagged proteasomes in HCT116 cells, we captured and structurally localized a set of transient, sub-stoichiometric proteasome-associated proteins. For each of these, we encountered substantial challenges when using only cryo-EM for structural analyses. A key innovation of this study is the use of high-fidelity in situ photo-crosslinking to preserve labile interactions prior to proteasome purification and capture distance information by identifying the crosslinked residues. Several known regulators, including UBE3C, were significantly enriched only in crosslinked samples. Combining in situ-derived crosslinks, we were able to filter AlphaFold predictions to generate a model for how UBE3C interacts with proteasomes, from its N-terminal end at RPN3 as it spans across RPN2 and RPN10 placing its more dynamic HECT domain near the RPN11 active site. From this position, UBE3C can readily ubiquitinate substrates that are stalled or degraded slowly to prevent premature substrate release. This positioning also suggests that UBE3C can ubiquitinate substrates near RPN11, which can then remove the chains en bloc^49^ with coupled translocation into the ATPase pore^50^. This proposed coordination of UBE3C and RPN11 activities provides a rationale for the UBE3C requirement for processive degradation of substrates by proteasomes^30,31,48^.

UBE3C is a therapeutic target for melanoma^61^, glioma^62^, renal cell carcinoma^63^, breast cancer^64^, cystic fibrosis^65^, and neurological disorders. The UBE3C paralog UBE3A is also present at proteasomes, where it binds to the extreme C-terminus of RPN10^12^. Its binding at this location suggests that UBE3A and UBE3C may be able to bind to proteasomes simultaneously; however, they do not seem to be able to substitute for each other functionally. Loss-of-function UBE3A mutations drive Angelman syndrome ^66–68^ while its elevated gene dosage correlates with autism spectrum disorders ^69^. Like UBE3A, biallelic loss-of-function mutations in UBE3C associate with Angelman-like symptoms^70^. In addition, a UBE3C intergenic fusion that loses the HECT domain is linked to distal hereditary motor neuropathy^71^. The informational provided here on UBE3C positioning at proteasomes can be used to guide future studies aimed at dissecting whether its role in these diseases relies on its activity at proteasomes.

We were able to resolve TXNL1 and PSMD5 at proteasomes by cryo-EM but their intrinsic dynamics posed challenges. The TXNL1 PITH domain was readily observed in our cryo-EM dataset at the base-lid interface near RPN2, RPN10, and RPN11. Its Trx domain however yielded weaker density and the in situ-derived crosslink between RPN13 Pru and TXNL1 provided confidence in its positioning at RPN2, near where RPN13 binds. TXNL1 is a thioredoxin-family protein previously implicated in redox regulation and protein quality control^57^. Its position is at a critical location in the substrate processing pathway, where deubiquitination and substrate commitment occur. This localization places TXNL1 where it could readily reduce oxidized substrates prior to their translocation into the CP – thus yielding peptides free of oxidative modifications. The functional role of proteasome-generated peptides is an active area of research, with recently discovered importance in microbial defense^18^ and neuron signaling^72,73^.

PSMD5 is an understudied proteasome interactor that promotes RP assembly^58^ and inhibits proteasome activity^59,60^. PhIX-MS revealed that PSMD5 interacts with all six RPT subunits, not just RPT1 and RPT2 as previously reported^59^, and these crosslinking data helped us resolved the structure of PSMD5-bound RP. With its C-terminus inserted into the central pore of the ATPase hexamer, PSMD5 weakens RPT interleaving to generate a unique, flatter ATPase structure. Co-fractionation MS suggests that PSMD5 is likely bound to most free RP particles in HCT116 cells and our structure suggests that PSMD5 interactions with RPT subunits stabilizes the ATPase ring until and following full assembly of the RP.

PSMD5 overexpression reduces 26S/30S proteasome assembly, supporting a model in which PSMD5 modulates base maturation and acts as a 26S/30S proteasome assembly checkpoint. This finding is consistent with its known downregulation in cancer^60^, as these cells increase proteasome activity to adapt to increased proteotoxic stress and reduction. PSMD5 may be fine-tuned to levels that enable RP assembly with seamless release and generation of 26S/30S proteasomes. Inhibitors of the proteasome CP are standard-of-care for hematological cancers^4^ and knowledge of how PSMD5 binds to the RP can be used to develop new therapeutic strategies for proteasome inhibition. For example, the C-terminal sequence that inserts into the ATPase ring can serve as a starting molecule for virtual screening of small molecules and/or peptide derivatives that block 26S/30S proteasome formation (Figure 6J).

In conclusion, we developed an approach to trap the conformational and compositional ensemble of protein complexes in cells with high-fidelity, and to use the crosslinked residue pairs as distance information to complement cryo-EM and other structural approaches. In this study, we provide a structural map of the proteasome interactome in intact human cells, revealing detailed information on how UBE3C, TXNL1, and PSMD5 engage proteasome complexes to regulate their assembly and promote their core activities. Our integrative approach establishes a generalizable framework for mapping dynamic protein assemblies in cells and offers new insights into the cellular logic of proteasome regulation by PSMD5 and activities of UBE3C and TXNL1.

## Limitations of the study

Photo-activated crosslinking is inefficient due to the high-propensity of the activated diazirine to quench with water^27^. This instability makes it difficult to detect the crosslinks and, in our study, crosslinked residue pairs were not detected for many crosslinked proteins that were enriched in the in situ crosslinked AP-MS experiments. As mass spectrometers improve for more sensitive detection, this limitation will be overcome. All experiments were performed in HCT116 cells under standard culture conditions; however, to prevent quenching of the crosslinker, the cells need to be exchanged into PBS (phosphate buffered saline) while incubating with SDA, which causes cellular stress. The proteasome interactome may differ across cell types, stress conditions, or developmental states. Further work will be needed to extend this strategy to diverse biological contexts.

## Resource availability

## Lead Authors

Francis J. O’Reilly; Kylie J. Walters

## Materials availability

Cell lines are available upon request.

## Data and code availability

The cryo-EM maps generated in this study have been deposited into the EMDB database with accession codes EMD-71740 (26S^TXNL1-1^), EMD-71741 (26S^TXNL1-2^), EMD-71810 (19S^TXNL1-1^), EMD-71737 (19S^TXNL1-2^), EMD-71791 (19S^ΔTXNL1-1^), EMD-71795 (19S^ΔTXNL1-2^), and EMD-71813 (19S^TXNL1-1, ΔRPT1, ΔRPT2^). The atomic models reported in this paper have been deposited into the PDB with accession codes 9PMO (26S^TXNL1-1^), 9PMQ (26S^TXNL1-2^), 9PRO (19S^TXNL1-1^), 9PMJ (19S^TXNL1-2^) and 9PRT (19S^TXNL1-1,ΔRPT1,ΔRPT2^). The raw micrographs have been deposited in EMPIAR with accession number EMPIAR-12891. All raw and processed Pulldown-ID mass-spectrometry datasets used in this study are available in the PRIDE database under accession code: Pulldown ID-1 (non-crosslinked WT vs RPN1 tag), PXD066281; Pulldown ID-2 (crosslinked WT vs RPN1 tag), PXD066280; Pulldown ID-3 (non-crosslinked WT vs RPN11 tag), PXD066279; Pulldown ID-4 (crosslinked WT vs RPN11 tag), PXD066278; Pulldown ID-5 (non-crosslinked WT vs PSMB4 tag), PXD066268; Pulldown ID-6 (crosslinked WT vs PSMB4 tag), PXD066284; and PSMD5 overexpression pulldown ID, PXD066535. All raw files and processed crosslinking MS datasets used in this study are available in the PRIDE database under accession code PXD066071, PXD066061, PXD066021 and PXD066073, and co-fractionation MS dataset under PXD066287. The AlphaFold3 model containing UBE3C, RPN2, RPN3, RPN8, RPN9, RPN10, RPN11, and RPN12 is available from ModelArchive under accession code ma-gxnsh.

Our AlphaFold2 prediction analysis Python pipeline is available on GitHub (https://github.com/katerinaatallahyunes/Proteasome-Interactome-AF-Predictions). All other data are available from the corresponding authors upon reasonable request.

## Acknowledgements

This research was supported by the Intramural Research Programs of the National Cancer Institute (NCI) of the National Institutes of Health (NIH, 1 ZIA BC011490 and 1 ZIA BC011627 to KJW and 1ZIA BC012114 to FJO), as well as federal funds from the NCI, NIH, under Contract No. HHSN26120150003I. The contributions of the NIH authors were made as part of their official duties as NIH federal employees, are in compliance with agency policy requirements, and are considered Works of the United States Government. However, the findings and conclusions presented in this paper are those of the authors and do not necessarily reflect the views of the NIH or the U.S. Department of Health and Human Services. We thank Ronald J. Holewinski (Laboratory Cancer Research Technology Program, Frederick National Laboratory for Cancer Research) for aiding acquisition of the co-fractionation MS dataset. This study utilized the Center for Structural Biology in the Center for Cancer Research (NCI, NIH) cryo-EM and NMR facilities and Biophysics Resource, the NCI Genome Modification Core and FACS facilities, and the Frederick Research Computing Environment (FRCE). We are grateful to Ines Chen and Andrea Graziadei for critical feedback on the manuscript.

## Author Contributions

K.J.W. and F.J.O. conceived of the project. K.L., K.A.-Y., and A.M.C. performed and analyzed all mass spectrometry experiments. K.A.-Y. performed the AlphaFold predictions and analysis. K.L. and H.N. characterized the edited cell lines and H.N. performed the biochemical assays on the purified samples and together with X.C. generated the sample and grids for cryo-EM data collection. X.C. and H.N. acquired the cryo-EM dataset and X.C. analyzed the data. X.C., H.N. and S.T. performed and analyzed all NMR experiments. M.G. and R.C. performed the CRISPR gene editing. R.E.C. and H.N. performed the binding assays involving UBE3C. S.T. and S.G.T. obtained and analyzed the ITC data. K.J.W. and F.J.O. supervised the project. All authors contributed to writing the manuscript.

## STAR Methods

### Design of CRISPR/Cas9 and donor plasmids

Guide RNAs (gRNAs) were designed in the region encompassing the final protein coding exon and 3′UTR using *sgRNA Scorer 2.0*^74^ (Table S1) and tested for cutting activity in 293T cells as previously described^32^. The chromosomal location and gRNA-binding regions within *PSMD14 and PSMB4* are listed in Table S2. Oligonucleotides corresponding to candidates IVT-2995, IVT-2997, JT-IVT-44 and JT-IVT-48, which were chosen for subsequent knock in experiments and named RPN11-01, RPN11-02, PSMB4-01 and PSMB4-02, respectively, were phosphorylated, annealed, and cloned into the pDG458 backbone using golden gate assembly (Table S3)^32^. pDG458 was a gift from Paul Thomas (Cat. No. 100900; Addgene plasmid).

A plasmid donor construct for HDR-based knock-in was generated using a combination of synthesized DNA (Twist Bioscience) of ∼800 bp of homology sequence 5′ and 3′ of the point of insertion and DNA sequence encoding TEV-biotin-P2A-mScarlet generated by PCR from existing plasmid. These DNA fragments were then cloned into the pGMC00018 vector (Cat. No. 195320; Addgene plasmid) using two sequential isothermal assembly cloning reactions^32^. The mScarlet in the donor plasmid was used as a fluorogenic selection markers with cell sorting by FACS.

### Expression plasmids

A plasmid for expressing the TXNL1 Trx domain (1-112 residues) in the pGEX6P1 vector with a GST tag at its N-terminus followed by PreScission protease cleavage site was synthesized by GenScript Biotech. To express GFP-tagged UBE3C, UBE3C x1 - x84 was in-frame at the N-terminal end of GFP in the pcDNA3.1 vector and synthesized by GenScript Biotech. To generate HA-PSMD5, an HA tag was fused to the N-terminal end of PSMD5 in pcDNA3.1 by GenScript. RPN10^UIM1-2^ (203-310), RPN10^UIM1^(196 – 272), RPN2 peptide (940-953) and RPN13 DEUBAD (253-407) were cloned into pGEX6T1 with a GST tag at the N-terminus followed by a precision protease cleavage site. RPN13 Pru (1-150) was cloned into the pRSET vector with a 6X-His tag at the N-terminus followed by a PreScission protease cleavage site.

### Transfection into HCT116 cells

1.5 × 10^5^ HCT116 cells (P3) purchased from American Tissue Culture Collection (CCL-247) were reverse transfected in a 6-well plate (Cat. No. 3506; Costar) with 2 μg of Cas9 plasmid and 5 μg of donor plasmid (RPN11/PSMB4-TEV-Biotin-P2A-mScarlet) by Lipofectamine 3000 (Cat. No. L3000015; Thermo Fisher Scientific, Inc) with Opti-MEM reduced serum medium (Cat. No. 31985070; Life Technologies) according to the manufacturer’s instructions. Transfected cells were incubated at 37 °C with 5% CO_2_ humidity for 72 h and then single sorted by FACS as previously described^32^.

### PCR genotyping

PCR genotyping was used to confirm integration of the knock-in tag at the genomic level. Different sets of primers (Integrated DNA Technologies, Inc) were designed against different regions of *PSMD14 and PSMB4* as well as the knock-in tag (Table S4). PCR was performed using the Phusion Plus PCR Master mix (Cat. No. F631S; Thermo Fisher Scientific, Inc) and the PCR products were analyzed by Nanopore sequencing.

### In situ photo-crosslinking of CRISPR/Cas9 knock-in cell lines

CRISPR/Cas9 knock-in cells were cultured in 100 mm × 25 mm dishes containing McCoy’s 5A medium (Cat. No. 16600082; Life Technologies Corp.) supplemented with 10 % (v/v) fetal bovine serum (Cat. No A5256701; Thermo Fisher Scientific) at 37 °C, 5 % CO_2_. Succinimidyl 4,4′-azipentanoate (SDA) was prepared as a 50 mg mL⁻¹ stock in DMSO. Confluent monolayers were rinsed twice with PBS (pH 7.4) and incubated with 4.4 mM SDA (1 mg mL⁻¹ in PBS) for 25 min at room temperature in the dark. Excess reagent was quenched by adding Tris-HCl (pH 8.0) to a final concentration of 50 mM and incubating for 5 min. Cells were then washed twice with PBS, overlaid with 5 mL PBS to prevent drying, and irradiated with 365 nm UV light for 10 s using an LED Cube 100 IC chamber paired with an LED Spot 100 HP IC lamp (Hönle UV Technology). After irradiation, cells were scraped, pelleted (200 × g, 5 min, 25 °C), snap-frozen in liquid nitrogen, and stored at −80 °C for downstream analyses.

### Sample preparation for Pulldown ID Mass Spectrometry (MS)

CRISPR/Cas9 knock-in HCT116 cells (RPN1/RPN11/PSMB4-tagged) were each cultured in 100 mm × 25 mm dishes. For every biological replicate, three dishes per cell line were subjected to the in-situ photo-crosslinking protocol described above, while the remaining three dishes served as non-crosslinked controls. Cells were lysed by Dounce homogenization on ice for 30 min in lysis buffer (50 mM HEPES, pH 7.5, 100 mM NaCl, 1 % NP-40, 10 % glycerol, 5 mM ATP, 10 mM MgCl₂, protease-inhibitor cocktail; Universal Nuclease, Cat. No. 88700; Thermo Fisher) with gentle inversion every 5 min. Lysates were clarified by centrifugation at 20,000 × g for 30 min at 4 °C, and the supernatants were incubated for 2 hours at 4 °C with pre-equilibrated MagReSyn Streptavidin MS beads (Cat. No. MR-STP002; ReSyn Biosciences). Beads were washed with wash buffer (50 mM HEPES, pH 7.5, 100 mM NaCl, 10 % glycerol, 2 mM ATP, 5 mM MgCl₂) four times for 10 min at 4 °C with rotation and collected on a magnetic rack.

Bound proteins were denatured in 8 M urea, 100 mM ammonium bicarbonate, and reduced with 5 mM dithiothreitol for 30 min at 37 °C, alkylated with 10 mM iodoacetamide for 30 min in the dark, and digested with Lysyl Endopeptidase (enzyme-to-protein ratio 1:100) for 4 h at room temperature with shaking. The urea was then diluted to 1.5 M with 100 mM ammonium bicarbonate and peptides were further digested overnight with trypsin (1:50) at room temperature. Digested peptides were acidified to pH 3, desalted on C18 StageTips, and dried in a vacuum concentrator (Eppendorf) for subsequent LC-MS/MS analysis using data-dependent acquisition (DDA) for label-free quantification (LFQ) method.

### LFQ data processing

The built-in “LFQ-MBR” workflow in FragPipe v22.0 (https://github.com/Nesvilab/FragPipe) was used for data analysis^75^. Briefly, MSFragger (v4.1) performed a closed search with initial precursor and fragment mass tolerances set to 20 ppm. Following mass calibration, MSFragger automatically refined these to narrower tolerances. The enzyme was specified as strict trypsin, allowing up to two missed cleavages. Carbamidomethylation of cysteine (+57.0215 Da) was set as a fixed modification, while methionine oxidation (+15.9949 Da), protein *N*-terminal acetylation (+42.0106 Da), and SDA (and hydrolysed SDA) modifications on lysine residues (+82.04186, +100.05243, +110.04801 Da) were included as variable modifications. After the search, MSBooster was used to calculate deep-learning scores, followed by Percolator for PSM rescoring, and Philosopher for false discovery rate (FDR) estimation at the peptide-spectrum match (PSM) level^76^. For label-free quantification, IonQuant (v1.10.27) was used with default settings^77^. The mass tolerance was set to 10 ppm, and the retention time tolerance to 0.4 minutes. Match-between-runs and MaxLFQ intensity calculation were enabled. Additional parameters included “min scans” set to 3, “min isotopes” to 2, and “MaxLFQ min ions” to 2.

### Data analysis for Pulldown ID MS

Label-free quantitative proteomics tables (PulldownID MS) were analyzed with QProMS v2 – “Quantitative PROteomics Made Simple” (https://github.com/FabioBedin/QProMS). After removal of contaminants, decoys and “only-identified-by-site” entries, proteins were retained if they were quantified in ≥ 80 % of replicates in at least one experimental group. All intensities were log_2_-transformed and variance-stabilizing normalization (VSN) was applied to correct for systematic between-sample effects. Missing values were handled with QProMS’ “mixed” imputation strategy, which treats data missing at random (MAR) and missing not at random (MNAR) separately: MAR values are replaced by the mean of the observed replicates, whereas MNAR values are drawn from a down-shifted Gaussian distribution (down-shift = 1.8 SD; width = 0.3 SD). Differential protein enrichment between conditions was assessed with a two-tailed Welch’s t-test. P-values were adjusted for multiple testing with the Benjamini– Hochberg false-discovery-rate (FDR) procedure (“BH truncation” in QProMS). Proteins exhibiting an absolute log₂ fold-change ≥ 2 (i.e., ≥ 4-fold) and FDR-adjusted P < 0.05 were considered significant and visualized in volcano plots generated within QProMS.

All raw and processed Pulldown-ID mass-spectrometry datasets used in this study are available in the PRIDE database under accession code: Pulldown ID-1 (non-crosslinked WT vs RPN1 tag), PXD066281; Pulldown ID-2 (crosslinked WT vs RPN1 tag), PXD066280; Pulldown ID-3 (non-crosslinked WT vs RPN11 tag), PXD066279; Pulldown ID-4 (crosslinked WT vs RPN11 tag), PXD066278; Pulldown ID-5 (non-crosslinked WT vs β4 tag), PXD066268; and Pulldown ID-6 (crosslinked WT vs β4 tag), PXD066284.

### Sample preparation for co-fractionation MS

Purified proteasome complexes were separated by size-exclusion chromatography on tandem Yarra SEC-4000 columns (3 µm, 300 × 7.8 mm; Phenomenex) equilibrated with 50 mM HEPES (pH 7.5), 100 mM NaCl, 2 % (v/v) glycerol, 1 mM ATP and 2 mM MgCl₂. A 200 µL sample was injected at a flow rate of 0.4 mL/min. Fractions (200 µL) eluting between 11 mL and 19.8 mL were precipitated with ice-cold acetone (−20 °C) and digested sequentially, first with Lys-C and then with trypsin. Peptides were desalted and cleaned using C18 StageTips as previously described. The sample from each fraction was further processed to LC-MS/MS.

### Data-dependent acquisition (DDA) for Label-Free-Quantification (LFQ)

Dried peptides were resuspended in 30 μL of 1.6 % acetonitrile with 0.1 % formic acid, vortexed and sonicated for 1 min before injecting 1 μg of estimated peptide sample onto an Orbitrap Eclipse Mass Spectrometer (Thermo Scientific) coupled to a Vanquish Neo HPLC system. Peptides were ionized using an EASY-Spray source and eluted over an EASY-Spray PepMap Neo 75 μm × 500 mm C18 column with LC–MS quality water or acetonitrile with 0.1% formic acid (mobile A and B, respectively). The flow rate was 0.25 μL/min using a gradient ranging from 1.6 % mobile phase B to 45 % mobile phase B.

Peptides were analyzed using the following MS global parameters: method duration of 120 min; infusion mode, liquid chromatography; expected LC peak widths, 30 s; advanced peak determination checked; default charge state of 2; EASY-IC internal mass calibration; NSI ion source; static spray voltage at 2,000 V in positive mode; static gas mode with a sweep gas setting of 2; ITT temperature of 280 °C. Samples were collected using the following shared scan parameters: Duty cycle of 3 s; MS-OT at 120,000 resolution; normal mass range; quadrupole isolation checked; scan range of 400–1,600; RF lens of 35%; a standard AGC target with auto injection time; one microscan in profile mode at positive polarity; EASY-IC checked; subbranch MIPS, peptide; subbranch intensity, 2.5 × 104; subbranch charge state 2–7; subbranch dynamic exclusion of one time after 30 s with a mass tolerance of 5 ppm; exclude within cycle checked; subbranch ddMS2 OT, isolation mode, quadrupole with a 1.4 m/z window; HCD with a fixed, normalized collision energy of 29; 15,000 resolution, normal mass range; auto scan range mode; standard AGC target; one microscan; centroid data.

### Purification of 26S proteasomes for PhIX-MS

Crosslinked cells were lysed with lysis buffer (50 mM HEPES pH 7.5, 100 mM NaCl, 1% NP-40, 10% glycerol, 5 mM ATP, and 10 mM MgCl2, supplemented with protease inhibitor cocktail and the universal nuclease (Cat. No. 88700; ThermoFisher Scientific, Inc)) using a Dounce homogenizer and incubated on ice for 15 min. The lysed sample was centrifuged at 20,000 x g for 30 min in a prechilled centrifuge at 4 °C. The supernatant was incubated for 2 h at 4 °C with High Capacity Neutravidin Agarose resin (Cat. No. 29202; Thermo Fisher Scientific, Inc) that was preequilibrated with lysis buffer. The resins were next washed with wash buffer (50 mM HEPES pH 7.5, 100 mM NaCl, 10 % glycerol, 1 mM DTT, 2 mM ATP, and 5 mM MgCl_2_) and then incubated for 2 h at 25 °C with TEV-digestion buffer (50 mM HEPES pH 7.5, 100 mM NaCl, 10 % glycerol, 1 mM DTT, 2 mM ATP, and 5 mM MgCl_2_) containing His-tagged TEV protease (Cat. No. 12575015; Thermo Fisher Scientific, Inc). To remove TEV enzyme, TALON Superflow resin (Cat. No. 28957502; Cytiva), pre-equilibrated with TEV-digestion buffer, was added to the mixture. The unbound mixture, containing cleaved 26S proteasome, was collected and protein was precipitated with ice-cold acetone.

### Offline peptide fractionation

For crosslinked peptide enrichment, peptides were fractionated on an ÄKTA Pure system (GE Healthcare) using a Superdex 30 Increase 3.2/300 (GE Healthcare) at a flow rate of 10 μL/min using 30 % (v/v) acetonitrile and 0.1 % (v/v) trifluoroacetic acid as the mobile phase at 4 °C. 50 μL fractions were collected from the elution volume 1.00 mL to 1.45 mL and dried for subsequent peptide strong cation exchange (SCX) chromatography or liquid chromatography–tandem mass spectrometry (LC–MS/MS) analysis.

SCX chromatography was performed on the same ÄKTA Pure system with a PolySULFOETHYL A™ column (100 mm × 2.1 mm, 3 µm, 300 Å; PolyLC Inc.) at 150 µL/min. Mobile phase A consisted of 10 mM KH₂PO₄ (pH 3.0) with 30 % (v/v) acetonitrile; mobile phase B was identical but contained an additional 1 M KCl. After sample loading, the column was washed with 3.5 % B, followed by a linear gradient to 40 % B to elute cross-linked peptides. Fractions of 200 µL were collected, partially dried (to remove acetonitrile), and cleaned on C18 StageTips before being analyzed via LC–MS/MS.

### LC-MS/MS analysis of crosslinking mass spectrometry

Peptides were analyzed using mass spectrometry global parameters as previously described^78^. High-field asymmetric waveform ion mobility spectrometry (FAIMS) was optionally applied depending on sample injection availability, using compensation voltages of −55, −45/−65, and/or −40/−70 at standard resolution. Targeted precursor selection ranges were also optionally used based on injection number and included the following m/z windows: 380–655, 500–755, 650–905, 750–1005, 900–1800, and 1000–1800. The BoxCar acquisition method was used with a 5.12-second duty cycle, targeted SIM scans acquired at 240,000 resolution, multiplex isolation of 10 ions (user-defined groups), custom AGC target with automatic injection time, and source fragmentation energy set to 10 V. Two BoxCar mass lists were used specifying center m/z values and isolation windows. Box 1: 414.1/30.2, 462.2/24.8, 504.1/23.6, 545.35/23.1, 587.8/25, 634.05/26.7, 686/31.4, 784.45/38.9, 829.65/48.5, 957.2/91.6. Box 2: 439.5/26.6, 483.45/23.7, 524.85/23.9, 566.15/24.3, 610.5/26.4, 658.85/28.9, 715.35/33.3, 786.65/43.5, 882.65/63.5, 1101/202. The AGC target for BoxCar acquisition was set to 50%.

### Data analysis of PhIX-MS

A recalibration to control for detector error was conducted on MS1 and MS2 based on the median mass-shift of high-confidence (<1% FDR) linear peptide identifications from each raw file. To identify crosslinked peptides, the recalibrated peak lists were searched against the forward (target) and the reversed sequences (as decoys) of crosslinked peptides using the Xi software suite (v.1.8.6; https://github.com/Rappsilber-Laboratory/XiSearch)^79^. The following parameters were applied for the search: MS1 accuracy = 2 ppm; MS2 accuracy = 5 ppm; enzyme = trypsin allowing up to 3 missed cleavages and 2 missing monoisotopic peaks; crosslinker = SDA with an assumed NHS-ester reaction specificity for K, Y, S, T, and protein *N*-termini; diazirine reaction specificity for A, C, D, E, G, H, I, K, L, P, S, T, V, Y, and protein *C*- and *N*-termini; fixed modifications = carbamidomethylation on cysteine (Ccm); variable modifications = acetylation on lysine and protein *N*-termini, oxidation on methionine, hydrolyzed SDA on lysines and protein *N*-termini. MS cleavage of SDA crosslinks was considered during searches.

Before estimating the false-discovery rate (FDR), the resulting matches were filtered to those having greater than two fragments matched with a non-cleaved SDA and at least five matches total per peptide. These candidates were then filtered to a 2 % residue-pair target-decoy false discovery rate (FDR), with an additional threshold limiting the protein-protein FDR to 5 % using XiFDR, boosting for PPIs (v.2.3.2)^80^.

In the RPN1 (datasets 1 and 2) pulldown, we identified 3,213 unique residue pairs (1,010 inter-protein). The RPN11 (dataset 3) pulldown yielded 3,371 unique residue pairs (1,847 inter-protein) at the same FDR threshold. When combined, the two datasets included 5,459 non-redundant crosslinked residue pairs. This includes 2,236 hetero-protein links describing 276 PPIs, with a combined estimated PPI-FDR of 7.7 % (Extended Data 3). In the PSMB4 (dataset 4) pulldown, we identified 1,798 unique residue pairs (955 inter-protein) at the same FDR threshold (Extended Data 3).

The crosslinking MS datasets used in this study are available in the PRIDE database under accession code PXD066071, PXD066061, PXD066021 and PXD066073.

### AlphaFold structure prediction

We predicted pairwise structural models between the 19 established RP subunits and the 22 proteins identified as directly crosslinked to the RP (Extended Data 5). Predictions were run using AlphaFold2-Multimer (v.2.3.2) with the following settings: 5 models, 3 recycles, templating enabled, and dropout disabled^81^. All predictions were performed using a local installation of AlphaFold2-Multimer on a Linux server using NVIDIA V100 GPUs.

We built a Python pipeline to automatically extract spatial, confidence, and accuracy metrics, enabling systematic evaluation of predicted protein-protein interactions. Model quality was primarily assessed using predicted TM-score (pTM), which reflects confidence in the structure of individual chains, and interface predicted TM-score (ipTM), which indicates confidence in the predicted inter-chain interface. We used these two scores to evaluate each prediction’s model confidence, which is 0.8*ipTM + 0.2*pTM. Predictions with an average confidence score above 0.65 (across all five models) were considered high confidence.

An AlphaFold3 prediction containing UBE3C, RPN2, RPN3, RPN8, RPN9, RPN10, RPN11, and RPN12 was run via AlphaFold Server^82^ with the default parameters.

### Purification of 26S proteasomes for cryo-EM

A frozen RPN1-tagged cell pellet (∼2 g) from twenty 145 mm x 20 mm cell culture dishes (Cat. No. 639160; Greiner Bio-one Inc.) was homogenized using a Dounce homogenizer (Cat. No. 1234F35; Thomas Scientific Inc.) in 12 mL of lysis buffer (Buffer 1: 50 mM Tris, pH 7.5, 10 % glycerol, 2 mM ATP, 5 mM MgCl_2_ and 1 mM DTT supplemented with Protease inhibitor tablet) followed by 7 freeze-thaw cycles. The lysate was then centrifuged at 17,000 x g for 15 min at 4 °C before aliquoting the supernatant, 1 mL each, in 1.5 mL Eppendorf tubes containing 100 μL of buffer equilibrated neutravidin agarose beads (Cat. No. 29204; Thermo Scientific). The bead: supernatant mixture was incubated for 2 h at 4 °C before washing with wash buffer (Buffer 2: 50 mM Tris, pH 7.5, 10 % glycerol, 2 mM ATP, 5 mM MgCl_2_ and 1 mM DTT) for 10 min at 4 °C. The washing step was repeated three times before incubating each samples in 500 mL TEV cleavage buffer (Buffer 3: 50 mM Tris, pH 7.5, 10 % glycerol, 5 mM ATP, 5 mM MgCl_2_, 1 mM DTT and 15 U mL^-1^ of AcTEV (Cat. No. 12575015; Life Technologies Inc.)) for 2 h at 25 °C. After the cleavage the solution containing the cleaved proteasome was separated from beads via centrifugation at 800 x g for 5 min at 4 °C. The cleaved proteasome complex was then concentrated using an Amicon filter with 100 kDa molecular weight cut-off (Cat. No. UFC510096; Millipore) to a final concentration of 0.75 mg/mL. The purified proteasome was aliquoted, flash frozen in liquid nitrogen, and stored in −80 °C.

### Cryo-EM sample preparation, grid preparation and data acquisition

To prepare cryo-grids, the purified proteasome samples were thawed on ice for 10-15 min and buffer exchanged with Buffer 4 (50 mM Tris, pH 7.5, 50 mM NaCl, 1.5 mM ATP-γ-S, 5 mM MgCl_2_ and 2 mM DTT) using a Zeba Micro Spin Desalting Columns (7k MWCO, Cat. No. 89883; Thermo Fisher).

Quantifoil grids (R 1.2/1.3 300 mesh, copper, Electron Microscopy Sciences) were glow discharged on each side for 30 s by using a Pelco easiGlow^TM^ glow discharge cleaning system at a negative discharge of 25 mA plasma current and 0.38 mbar residual air pressure. The grids were plunge-frozen by using a Leica EM GP2 cryoplunger (Leica Microsystems, Germany) operated at 22 °C and 80 % relative humidity. For each grid, 2.5 μL of proteasome sample (0.75 mg/mL) was applied to the carbon side and 1.0 μL of proteasome sample (0.75 mg/mL) was applied to the non-carbon side. The grids were blotted for 2, 2.5, 3, and 4 s at their non-carbon side and vitrified by plunge freezing into liquid ethane cooled by liquid nitrogen. The plunge-frozen grids were then clipped and stored in cryo-grid boxes under liquid nitrogen until the data collection step.

Cryo-EM data was acquired by using a Talos Arctica G2 electron microscope (Thermo Fisher Scientific) equipped with a K3 direct electron detector (Gatan) and an energy filter, operating at 200 kV in super resolution mode (pixel size 0.405 Å/pixel, × 100,000 nominal magnification). 40 frames per movie were acquired for a total dose of approximately 55.6 electrons per Å^2^ and an exposure time of 2.5 s. Data were collected using the EPU program (Thermo Fisher Scientific), with defocus values ranging from −2.0 to −0.8 μm. A total of 27,259 movies were collected from a selected cryo-grid.

### Cryo-EM image processing

All cryo-EM data processing was performed by using cryoSPARC 4.5.3^83^. A flowchart of the data processing is displayed in Figure S5A and S5B, and a summary of cryoEM reconstruction statistics is listed in Table S5 and S6. 27,259 dose-fractionated movies were gain-reference corrected, aligned, dose-weighted and summed to single-frame micrographs by patch motion correction, after which constant transfer function (CTF) estimation was done by patch CTF estimation. The micrographs were visually inspected and those with broken or thick ice, or CTF resolution > 5 Å were excluded from the stack, leaving 26,404 micrographs. An initial set of 350,329 particles were blob picked and particles were extracted from 2,330 micrographs, with a box size of 756 pixels and binned by 2x. Templates were created by selecting 9 classes by running 2D classification and template picking identified 254,030 particles from 2,330 micrographs and new templates were created by selecting 46 classes followed by 2D classification. An additional template picked from 6,772 micrographs identified 792,960 particles, for which 2D classifications were performed and new templates created by selecting 23 classes with 184,938 particles. A total of 5,299,947 particles were picked from 26,404 micrographs by template picking and extracted by using a box size of 410 pixels.

Particle stacks were subjected to one round of ab initio model generation and 3D heterogeneous refinement with binning into 3 classes. One class of 1,883,666 particles showed CP density with partial RP density capped on each side (Class 1 in Figure S5A) and was further subjected to one round of ab initio model generation and 3D heterogeneous refinement to remove unfolded particles. 1,199,862 particles were subjected to a heterogeneous refinement into 3 classes, and each class showed RP density on one side or both sides of an CP. Particles from each class were re-extracted by using a box size of 756 pixels, followed by non-uniform refinement and local refinement. To separate particles into individual proteasome states, two masks for RP plus α ring were generated for each class and particles were subjected to alignment-free 3D classification (8 classes, filtered resolution of 15 Å, initialization mode PCA, initial structure low-pass resolution of 30 Å). A total of 48 classes were sorted, pooled based on the conformation of the RP, and particle subtraction applied to remove CP signal beyond the RP-bound α-ring. CTF refinement, reference-based motion correction, non-uniform refinement, and local refinement were next applied. In additional to the 3 classes (105,579 particles) identified as the proteasome ground state (S_A_) and 5 classes (257,779 particles) identified as open-gate states (S_D_), 15 classes with 425,582 particles show extra density near RPN2, RPN11 and Rpn10 (Figure S5A). The heterogeneity of these particles showing extra density was analyzed and clustered in cryoDRGN^84^ followed by re-extracting particles using a box size of 480 pixels. Next, particle subtraction and local refinement were done to yield 26S^TXNL1-1^ and 26S^TXNL1-2^ maps with overall resolutions of 3.82 Å (143,635 particles) and 4.0 Å (100,351 particles), respectively, as analyzed by the Gold Standard Fourier Shell Correlation (GSFSC) in cryoSPARC (Figure S5A, S5C-S5H).

Class 2 (1,911,783 particles) showed less RP density and no TXNL1 density (Figure S5A) and we therefore focused on class 3 (Figure S5B). After ab initio model generation and heterogeneous refinement (3 classes), one class of open-gate 26S proteasome (S_D_, 424,557 particles), one class of unfold particles (273,512), and one class of free RP (595,212) particles were identified (Figure S5B). Free RP particles were subjected to one more round of heterogeneous refinement, resulting in two classes of free RP particles and one class of RP some CP density visible. The two classes of free RP particles were re-extracted for particles by using a box size of 432 pixels and CTF refinement, reference-based motion correction, non-uniform refinement, and local refinement were performed to generate 19S^TXNL1-1,ΔRPT1,ΔRPT2^ and 19S**^Δ^**^TXNL1-1^ maps with overall resolutions of 3.02 Å (228,010 particles) and 3.92 Å (102,897 particles), respectively, as judged by the GSFSC in cryoSPARC (Figure S5B, SI-SN). An alignment-free 3D classification masked on the RPT1-RPT2-RPT5-RPT6 AAA+ domain and RPN1 (6 classes, filtered resolution of 15 Å, initialization mode PCA, initial structure low-pass resolution of 30 Å) of 19S^TXNL1-1,ΔRPT1,ΔRPT2^ resulted in a combined 3 classes of particles (127,238) with extra density adjacent to RPN1-RPN10-RPN11 or RPT1/RPT2 (Figure S5B). These particles were then subjected to focused local refinement and 3D variability analysis and clustering to generate the 19S^TXNL1-1^ map with an overall resolution of 4.58 Å (34,566 particles, Figure S5O-S5Q). A parallel process was applied to 19S**^Δ^**^TXNL1-1^ to generate the 19S**^Δ^**^TXNL1-2^ map with an overall resolution of 4.24 Å (35,258 particles, Figure S5R-S5T). Heterogeneous refinement of the class that showed RP with less CP particle density separated the particles of free RP and 26S proteasomes. These free RP particles were then re-extracted by using a box size of 432 pixels and subjected to CTF refinement, reference-based motion correction, non-uniform refinement, and local refinement, generating the 19S^TXNL1-2^ map with an overall resolution of 4.22 Å (39,459 particles, Figure S5U-S5W).

The 26S^TXNL1-1^, 26S^TXNL1-2^, 19S^TXNL1-1,ΔRPT1,ΔRPT2^, 19S^TXNL1-1^ and 19S^TXNL1-2^ maps were then used for further model building and refinement. All GSFSC and viewing direction distribution plots were generated in cryoSPARC, and the local resolution estimation maps were generated in cryoSPARC or the Phenix suite^85^.

### Model building and structure analyses

RP and CP α subunits from a human 26S proteasome atomic model (PDB 7W3I) and the TXNL1 PITH domain structure (PDB 1WWY) were fitted as rigid bodies into the 26S^TXNL1-1^ and 26S^TXNL1-2^ density maps by using UCSF ChimeraX^86^. For the 26S^TXNL1-1^ density map, the TXNL1 Trx domain atomic model (PDB 1GH2) was also fitted as a rigid body into the density. Similarly, RP subunits from PDB 7W3I and the TXNL1 PITH domain from PDB 1WWY were fitted as rigid bodies into the 19S^TXNL1-1^ and 19S^TXNL1-2^ density maps by using UCSF ChimeraX^86^. The AlphaFold3 model of PSMD5 with the ATPase ring verified by crosslinking distances was used to fit PSMD5 into the extra density adjacent to RPT1/RPT2 in the 19S^TXNL1-1^ map. For 19S^TXNL1-1,ΔRPT1,^ **^Δ^**^RPT2^, the RP subunits except RPN1, RPT1, and RPT2 from a proteasome atomic model (PDB 7W3I) was fitted as a rigid body. To assist model interpretation, the refined maps were processed by EMready^87^. The amino acid rotamers and peptide bonds were flipped to increase Ramachandran favorability, decrease rotamer outliers, reduce clashes, and improve fit to the densities by using ISOLDE^88^ and Coot^89^. The structural models were further refined by iterative cycles of manual building in Coot^89^ and real-space refinement in the Phenix suite^85^. The geometry and real-space correlation validation were performed by using the phenix.validation_cryoem module in Phenix suite^85^. A summary of model building and validation statistics is listed in Table S5-S6. All low-pass-filtered maps were generated in CryoSPARC with volume tools^83^.

### Native gel electrophoresis and in-gel peptidase activity

Purified proteasome samples from RPN1-tagged, RPN11-tagged, and PSMB4-tagged cells were checked by in-gel peptidase activity assays following native-PAGE. 2 μg of each proteasome sample was loaded into a native-PAGE gel (3-8% Tris Acetate NuPAGE gel, Cat. No. EA0375; Thermo Fisher Scientific, lnc.) and run for 5.5 h at 150 V at 4°C. The gel was incubated for 15 min at 37 °C in in-gel activity assay buffer (50 mM Tris-HCl, pH 7.5, 2 mM ATP, 5 mM MgCl_2_,) with 12.5 μM suc-LLVY-AMC peptide (Cat. No. 10008119; Cayman Chemical Co, Inc.) and then imaged by a Gel Doc EZ Imager (Bio-Rad Laboratories, Inc.). The gel was next incubated again with in-gel activity buffer with addition of 0.02 % SDS and imaged again.

### Overexpression of UBE3C^N-term^-GFP and HA-PSMD5

A 10 cm culture dish was seeded with 1.1 million cells for 18 h, after which 5 μg of UBE3C^N-term^-GFP plasmid was added. For PSMD5 over-expression, RPN1-tagged cells were transfected in either 6-well plates (Cat. No. 3506; Costar) or 10 cm dishes using 1 µg or 5 µg of HA-PSMD5, respectively. All transfections were performed with Lipofectamine 3000 (Cat. No. L3000015; Thermo Fisher Scientific) in Opti-MEM reduced serum medium (Cat. No. 31985070; Life Technologies), following the manufacturer’s protocol. The cells were incubated at 37 °C in a humidified atmosphere containing 5 % CO₂ for 48 h before harvesting.

### Pull-down experiment and immunoblotting

The pull-down experiment for RPN1-tagged cells overexpressed without or with UBE3C^N-term^-GFP was performed using Dynabeads MyOne Streptavidin T1 beads (Cat. No. 65601; Life Technologies Corp.). The samples were lysed using lysis buffer (50 mM Tris, pH 7.5, 100 mM NaCl, 1 mM DTT, 5 mM ATP, 5 mM MgCl2, 10 % glycerol, 0.05 % NP-40 and supplemented with protease inhibitor cocktail) and then the lysate was centrifuged to remove cell debris. The total protein amount in the supernatant was measured by Pierce 660 nm protein assay reagent (Cat. No. 22660; Thermo Fisher Scientific) and 1 mg of total protein was incubated with streptavidin beads for 2 h followed by washing 2-3 times with wash buffer (50 mM Tris, pH 7.5, 100 mM NaCl, 1 mM DTT, 5 mM ATP, 5 mM MgCl_2_, and 10 % glycerol). The bound proteasome complex was eluted with addition of 2X SDS loading dye containing 50 mM DTT. Immunoblotting was done by overnight or for a 1 h incubation with primary antibodies (GFP (Cat. No. 2955S; CST), UBE3C (Cat. No. A304-123A; Bethyl Laboratories Inc.), PSMB5 (Cat. No. BML-PW8895; Enzo Life Sciences Inc.), β-actin (Cat. No. 4970T; CST), RPN11 (Cat. No. 4197S; CST), RPN1 (Cat. No. 25430S; CST)) in 5 % skim milk-Tris buffered saline with 0.1 % Tween 20 (TBST), followed by washing 3-4 times for 10 min each, and 1 h incubation with HRP-conjugated anti-rabbit (Cat. No. A16110; Life Techonologies) or anti-mouse (Cat. No. A9917; Sigma) secondary antibodies. The blots were washed again with TBST for 3-4 times (10 min each) before developing them on autoradiography films (Cat. No. 1141J52; Thomas Scientific, Inc) by addition of Pierce ECL chemiluminescent substrate (Cat. No. 32106; Thermo Fisher Scientific, Inc).

The pull-down experiment using RPN1-tagged cells, with or without HA-PSMD5 overexpression, was performed following the same procedure described for Pulldown ID mass spectrometry (MS) above. Immunoblotting was done by 1 h incubation with primary antibodies; HA-HRP (Cat. No. 2999S; CST), RPN1 (Cat. No. 25430S; CST), PSMB5 (Cat. No. 12919S; CST), ECM29 (Cat. No. AB28666; Abcam), β-actin (Cat. No. AB8226; Abcam) in Pierce™ Protein-Free T20 (TBS) Blocking Buffer (Cat. No. 37571; Thermo Fisher Scientific). After three 10-min washes in TBST, blots were incubated for 1 h with HRP-conjugated anti-rabbit (Cat. No. 31460; Life Techonologies) or anti-mouse (Cat. No. 31430; Life Techonologies) secondary antibodies, washed again, and developed on a ChemiDoc imaging system (Bio-Rad) using SuperSignal West Pico PLUS substrate (Cat. No. 34577; Thermo Fisher Scientific, Inc). The raw and processed mass-spectrometry datasets for RPN1 pulldowns with PSDM5 overexpression are available in the PRIDE database under accession code PXD066535.

### Chymotrypsin-like proteasome activity assay

Following transient overexpression, the harvested cells were resuspended in lysis buffer containing 50 mM HEPES (pH 7.5), 100 mM NaCl, 5 % glycerol, 0.1 mM EDTA, 2 mM ATP, and 4 mM MgCl₂, and lysed by repeated freeze–thaw cycles. Protein concentrations were determined using the Pierce™ Dilution-Free™ Rapid Gold BCA Protein Assay Kit (Thermo Fisher Scientific), according to the manufacturer’s instructions. Aliquots containing 50 µg of total protein were prepared and adjusted to a final volume of 50 µL with lysis buffer. Proteasome activity was measured using the Proteasome-Glo™ Chymotrypsin-Like Assay Kit (Promega), following the manufacturer’s protocol. Luminescence was detected using an Infinite® M200 PRO microplate reader (Tecan).

### Protein expression and purification

All the bacterial expression plasmids were expressed in *Escherichia coli* strain BL21(DE3) cells (Cat. no. EC0114; Thermo Fisher Scientific, lnc) expect for RPN13 Pru and GST-RPN2 (940-953), for which BL21(DE3) pLysS cells (Cat. No. C606003; Life Technologies Corp.) were used). The cells were grown at 37 °C to optical density at 600 nm of 0.5-0.6 and induced for protein expression by addition of 0.5 mM of isopropyl-1-thio-β-D-thiogalactopyranoside (IPTG, Cat. No. P101010; UBPBio). Post induction, the cells were grown at 17 °C for 18 h before harvesting them via centrifugation at 4,550 x g for 30 min at 4 °C. The cells were lysed by sonication, and cellular debris was removed by centrifugation at 31,000 x g for 30 min. The supernatant was incubated either with glutathione S-sepharose 4B resin (Cat. no. GE17075605; Millipore Sigma) or Talon Metal Affinity resin (Cat. No. 635502; Takara Bio Inc.) for 2 h. GST-RPN10^UIM1-2^, GST-RPN10^UIM1^ and GST-RPN2 (940-953) were eluted from the resin using 20 mM of reduced glutathione (Cat. No. G4251; Sigma) in 20 mM Tris (pH 7.5), 300 mM NaCl, and 10 mM β-mercaptoethanol (βME). TXNL1 Trx, RPN13 DEUBAD, and RPN13 Pru were eluted from the glutathione and Talon resin by overnight incubation with 50 units of PreScission protease (Cat. No. 27084301; Cytivia Life Sciences) in 50 mM Tis buffer (pH 7.5), 100 mM NaCl and 1 mM DTT. The eluted proteins were subjected to size exclusion chromatography with a HiLoad 16/600 Superdex 75 pg column on an FPLC system for further purification. ^15^N ammonium chloride was used for isotopic labeling of RPN13 Pru and RPN13 DEUBAD proteins.

### NMR experiment

Aliquots of ^15^N-labeled RPN13 Pru, ^15^N-labeled RPN13 DEUBAD, and unlabeled TXNL1 Trx were dialyzed using Slide-A-Lyzer mini dialysis devices into identical NMR buffer (20 mM sodium phosphate, 50 mM NaCl, pH 6.0) without any reducing reagent. For the oxidation of ^15^N-labeled RPN13 Pru and ^15^N-labeled RPN13 DEUBAD, the proteins were incubated with 0.05% H_2_O_2_ on ice for 10 min and the final round of buffer exchange into NMR buffer was done to remove remaining H_2_O_2_ from the samples. For the reduced ^15^N-labeled RPN13 Pru and ^15^N-labeled RPN13 DEUBAD samples, the purified proteins were buffer exchanged into NMR buffer containing 1 mM DTT.

2D ^1^H-^15^N HSQC spectra were recoded for 20 μM of ^15^N-labeled RPN13 Pru preincubated with equimolar unlabeled RPN2 peptide, and next with 5-fold, 10-fold, or 15-fold molar excess unlabeled TXNL1 Trx. ^1^H-^15^N HSQC was also recorded for 22 µM of reduced or oxidized ^15^N-labeled RPN13 Pru, and 22 µM of oxidized ^15^N-labeled RPN13 Pru with 4-fold excess of TXNL1 Trx (88 µM). ^1^H-^15^N HSQC spectra was also recorded for 25 µM of reduced or oxidized ^15^N-labeled RPN13 DEUBAD, and 25 µM of oxidized ^15^N- labeled RPN13 DEUBAD with 4-fold excess of TXNL1 Trx (100 µM). All of the NMR experiments were recorded at 10 °C on a Bruker Advance 600 MHz or 900 MHz spectrometer. The data were processed by NMRPipe^90^ (version 11.5) and spectra analyzed and visualized with XEASY^91^ or NMRFAM-SPARKY^92^ (version 3.190) by using NMRbox^93^.

### ITC experiment

For the ITC experiments, RPN13 Pru and TXNL1 Trx proteins were dialyzed utilizing a Slide-A-Lyzer mini dialysis device either in reducing ITC buffer (20 mM sodium phosphate, 50 mM NaCl, 10 mM βME, pH 6.0) or in non-reducing ITC buffer (20 mM sodium phosphate, 50 mM NaCl, pH 6.0). For the TXNL1 Trx-treated RPN13 Pru sample, RPN13 Pru was first oxidized as described above and then incubated with 3-fold molar excess of TXNL1 Trx for 2 h and then removed from the sample by size exclusion chromatography. All the ITC experiments were performed at 25 °C on a MicroCal iTC200 system (Malvern, PA, USA), with RPN13 Pru in the cell and RPN2 (940-953) in the syringe. One aliquot of 0.5 µL followed by 18 aliquots of 2.1 µL of 156 µM, 205.7 µM, or 160 µM RPN2 peptide was injected at 1,000 r.p.m. into the calorimeter cell (volume 200.7 µL), which contained either 15.6 µM (reduced), 20.57 µM (oxidized), or 16 µM (TXNL1 Trx-treated oxidized) RPN13 Pru in reducing or non-reducing conditions. Blank experiments were performed by replacing protein samples with buffer and the resulting data subtracted from the experimental data during analyses. The integrated interaction heat values were normalized as a function of protein concentration, and the data were fit with MicroCal Origin 7.0 software. Binding was assumed to be at one site to yield a binding affinity K_a_ (1/K_d_), stoichiometry and other thermodynamic parameters.

### UBE3C pull-down experiment

8 nmoles of GST, GST-RPN10^UIM1-2^ and GST-RPN10^UIM1^ were bound to 25 μL slurry of glutathione S-sepharose 4B resins for 1 hour at 4 °C. The beads were spun down (300 x g) and washed 2-3 times to remove the unbound excess proteins before incubating them with 1 mg of cell lysate from HCT116 cells, lysed in lysis buffer (50 mM TrisHCl (pH 7.5), 150 mM NaCl, 1 mM phenylmethylsulfonyl fluoride (PMSF, Cat. No. 10837091001; Sigma) and 1% Triton-X supplemented with EDTA-free protease inhibitor cocktail (Cat. No. A32955; Thermo Fisher Scientific, lnc.) for 3 h at 4 °C. The beads were again spun down and washed with lysis buffer before eluting the sample by adding 37.5 μL of 3X LDS loading dye to each sample. The samples were heated at 72 °C before loading them on an SDS gel and then transferred onto a PVDF membrane. Before immunoblotting with UBE3C antibodies, the PVDF membrane was stained with Ponceau stain to check the recombinant proteins amount on the membrane.

---

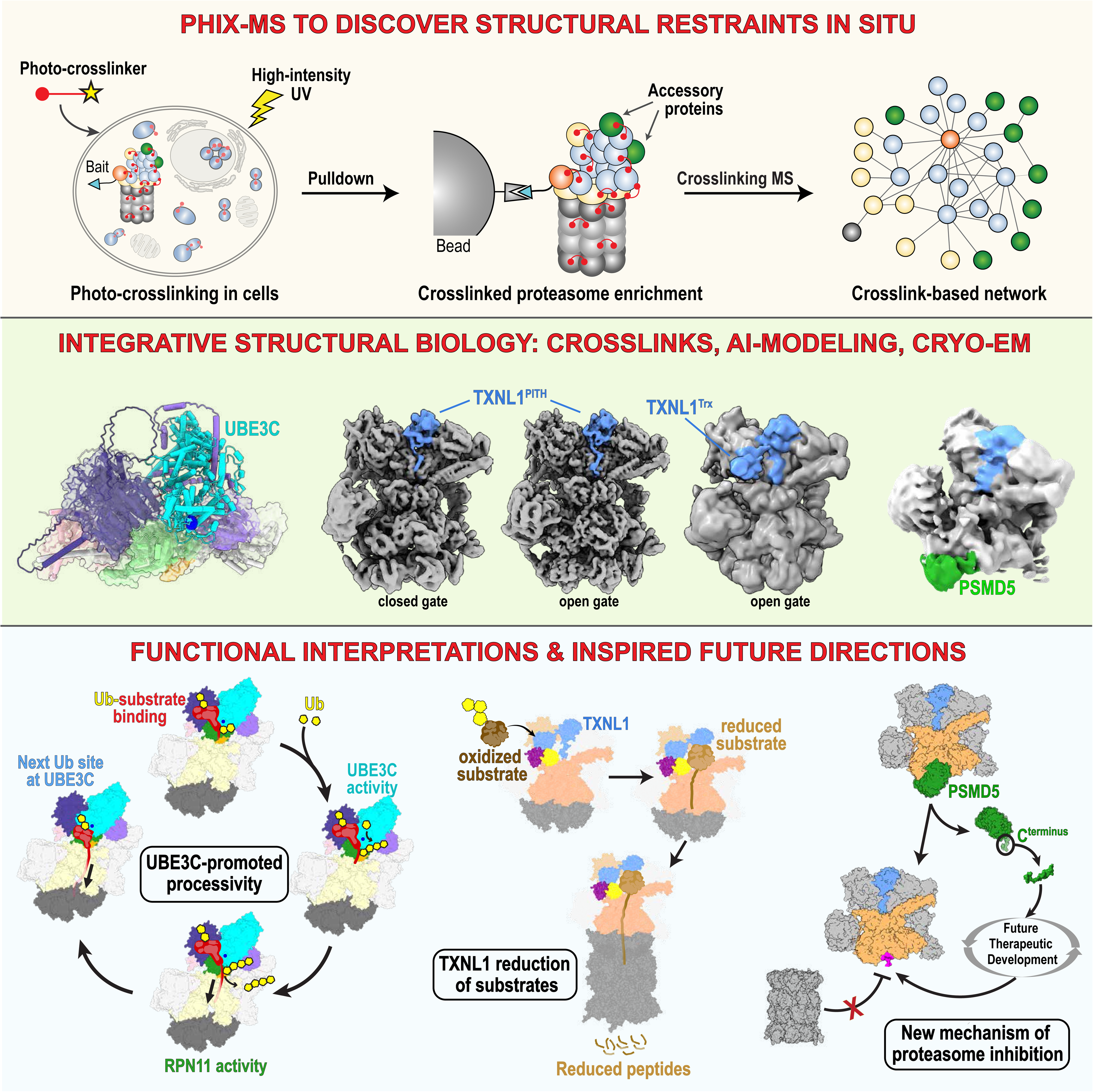

