## Supplementary Tables and Figures for "Structures of dynamic interactors at native proteasomes by PhIX-MS and cryoelectron microscopy"

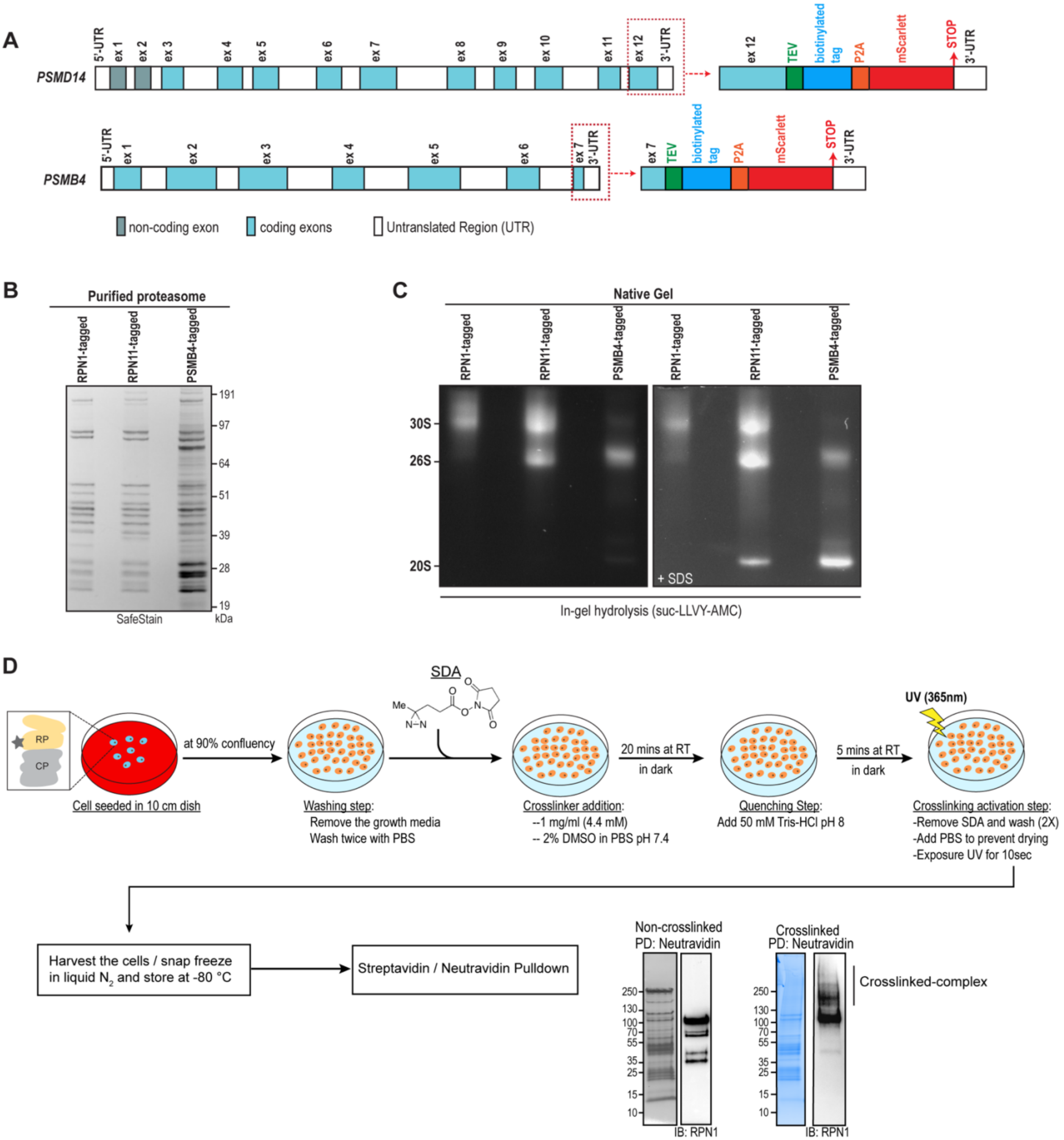

**Figure S1. Gene editing and crosslinking methodology for RPN1-, RPN11-, and PSMB4- tagged cell lines, related to Figure 1**. **(A)** Schematic representation of the *PSMD14* and *PSMB4* gene with their respective coding and non-coding exons and untranslated regions (UTR). CRISPR editing of *RPN11* and *PSMB4* at their terminal 3'-exon introduced a purification handle which contains a TEV-cleavable biotinylated tag for purification and P2A-mediated auto cleavable mScarlet signal. **(B)** Purified proteasome complexes (1 μg) from RPN1-, RPN11-, and

PSMB4-tagged cell lines were separately loaded onto a denaturing SDS-PAGE gel and stained with safe stain. **(C)** Purified proteasome complexes (2 μg) from RPN1-, RPN11-, and PSMB4-tagged cell lines were subjected to native-PAGE gel followed by in-gel peptidase activity by incubating with gel peptidase buffer (containing suc-LLVY-AMC peptide) without or with SDS. **(D)** Illustration of the in situ photo-crosslinking workflow. In short, tagged cells were grown to 90% confluency, media was removed via PBS washes, and cells were crosslinked with SDA (4.4 mM, 20 mins., room temperature) in the dark. Cells were re-washed with PBS and irradiated with 365 nm wavelength light for 10 seconds (see methods). Cells were then harvested, lysed, and the tagged proteins were enriched. Crosslinking was visualized as band streakiness and upward shifting on a gel via immunoblotting.

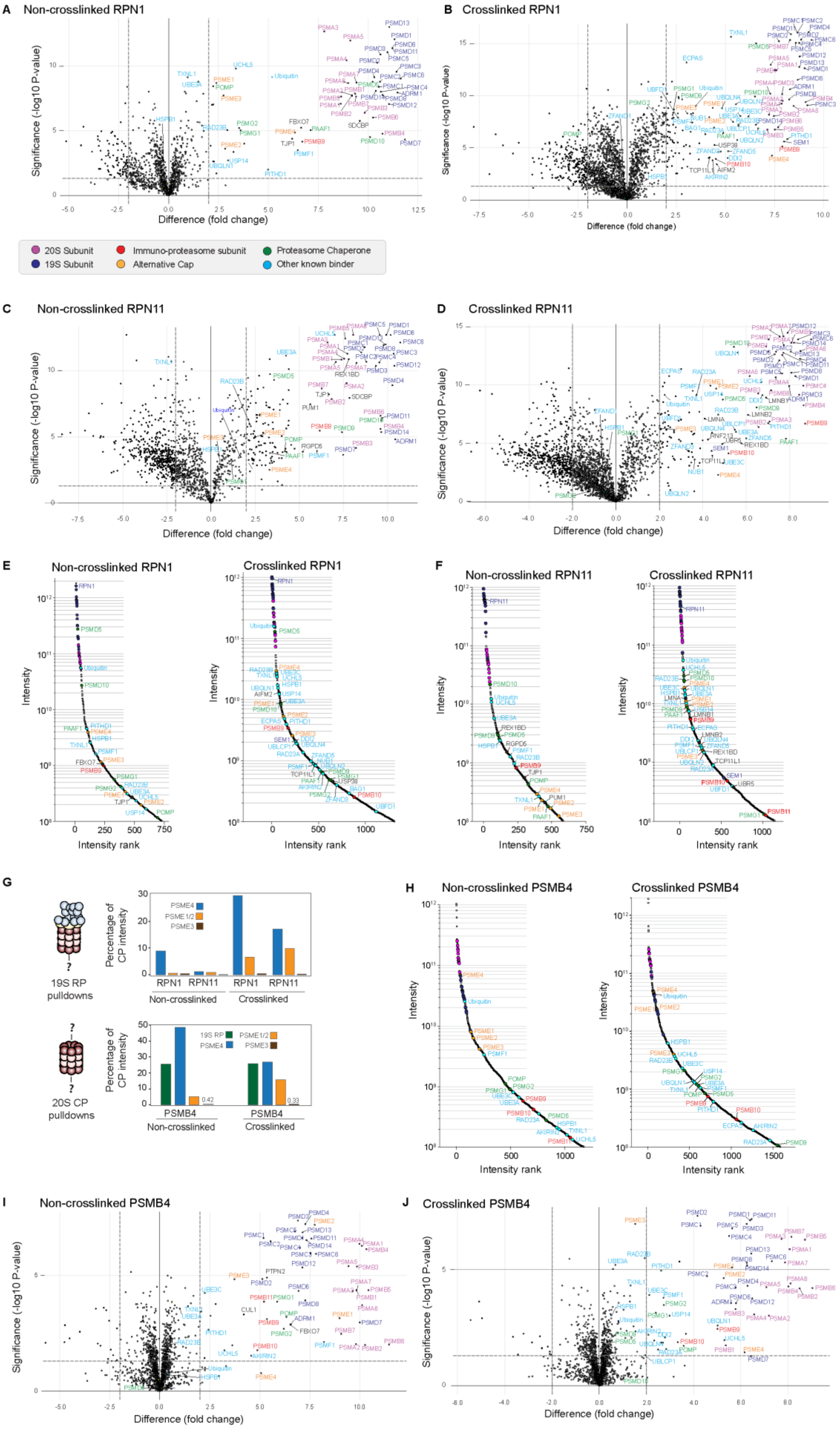

A Non-crosslinked RPN1
B Crosslinked RPN1
Significance (-log10 P-value)
Difference (fold change)
20S Subunit
19S Subunit
Immuno-proteasome subunit
Alternative Cap
Proteasome Chaperone
Other known binder
C Non-crosslinked RPN11
D Crosslinked RPN11
E Non-crosslinked RPN1
Crosslinked RPN1
F Non-crosslinked RPN11
Crosslinked RPN11
Intensity
Intensity rank
G
19S RP pulldowns
Percentage of CP intensity
PSME4
PSME1/2
PSME3
RPN1
RPN11
Non-crosslinked
Crosslinked
20S CP pulldowns
19S RP
PSMB4
0.42
0.33
H Non-crosslinked PSMB4
Crosslinked PSMB4
I Non-crosslinked PSMB4
J Crosslinked PSMB4

**Figure S2. AP-MS enrichment plots from biotin-tagged proteasome subunit pulldowns, related to Figure 1. (A-D, I-J)** Volcano plots depicting enrichment of pulled-down proteins for non-crosslinked (**A, C, I**) or crosslinked (**B, D, J**) for RPN1 (**A, B**), RPN11 (**C, D**), and PSMB4 (**I, J**) affinity-purification mass spectrometry (AP-MS) experiments. Enrichment is measured against pulldowns from parental HCT116 cells, either non-crosslinked (**A, C, I**) or crosslinked (**B, D, J**). Proteins are color-coded depending on whether they are part of the CP (pink), RP (dark blue), an immuno-proteasome subunit (red), an alternative cap (orange), a proteasome chaperone (green), or another known proteasome binder (cyan). The dashed line on the y-axis indicates the level of significance (P-value with BH correction < 0.05) and the dashed lines on the x-axis indicate the difference in abundance ($log_2$-fold change) between tagged and wild-type pulldown samples. Each dataset contains three independent biological replicates injected twice on the mass spectrometer. FragPipe/MSFragger was used to generate label-free quantification (LFQ) for the identified proteins, and that output was subsequently analyzed using QProMS to generate volcano plots, which were redrawn using custom Python scripts. **(E, F, H)** Ranked intensity plots for protein abundances using LFQ for non-crosslinked (left) and crosslinked (right) RPN1 **(E)**, RPN11 **(F)**, and PSMB4 **(H)** AP-MS experiments. **(G)** Bar plots for the non-crosslinked and crosslinked AP-MS datasets that compare the abundances of 19S RP (green), PSME4 (blue), PSME1/2 (orange), and PSME3 (brown) as the percentage of 20S CP intensity for 19S RP (top) or 20S CP pulldowns (bottom).

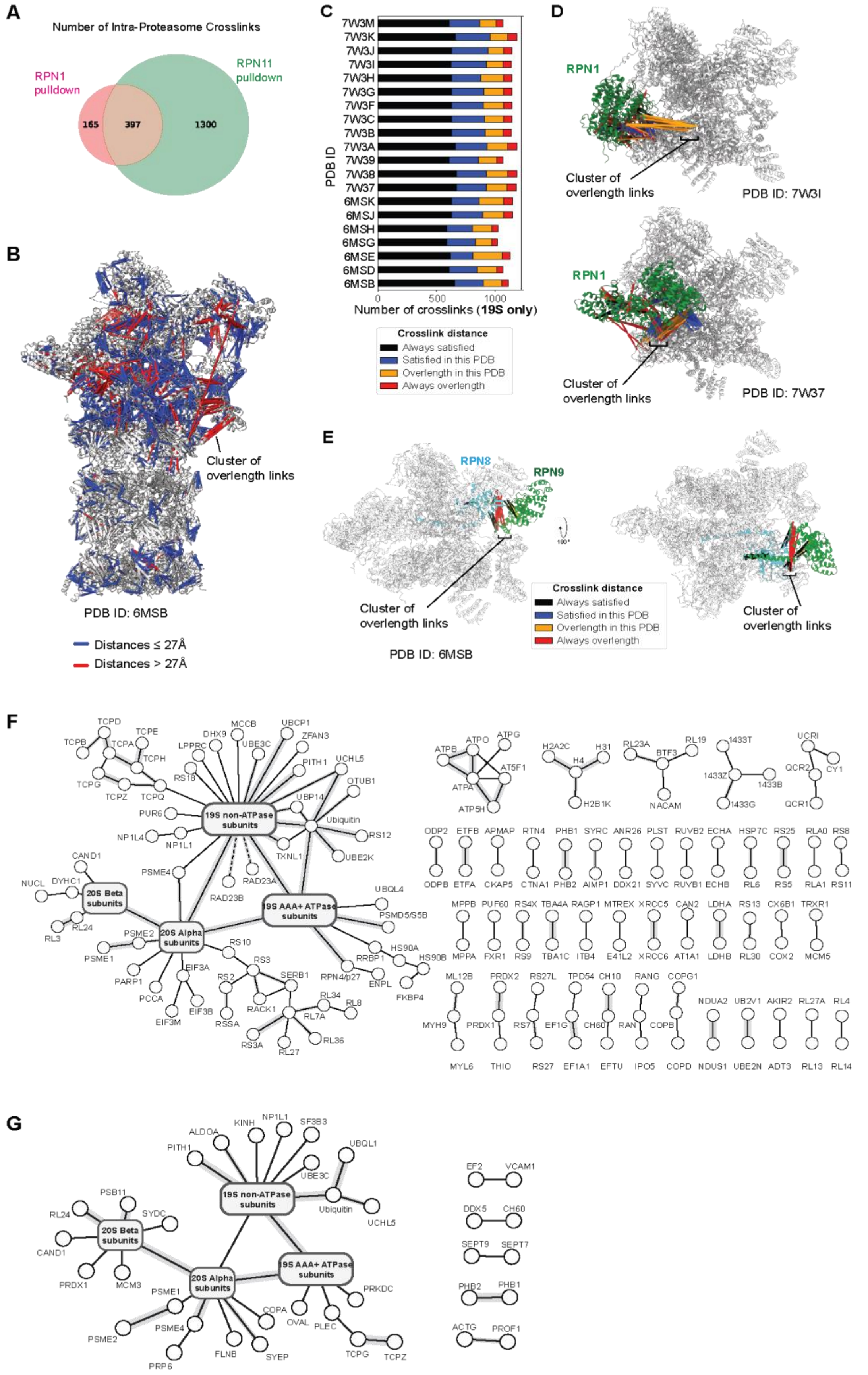

**Figure S3. Crosslinks on proteasome structures, related to Figure 2**. **(A)** Overlap of crosslinked residue pairs between subunits of the proteasome identified in the RPN1 and RPN11 pulldowns. Data come from 90 or

152 MS acquisitions from crosslinked RPN1 or RPN11 pulldown material, respectively. **(B)** A proteasome structure (PDB ID: 6MSB) with crosslinks mapped as pseudobonds and colored depending on Euclidean distance between residue-pair Cα–Cα distances. Crosslink-based distances of $\leq$ 27 Å are considered 'satisfied' (blue). Crosslinks that exceed this distance are colored in red. **(C)** Bar plot illustrating 20 different proteasome models (y-axis, PDB IDs) and whether the mapped crosslinks (x-axis, number of crosslinks mapped to the 19S subunit) satisfied the distance restraint. 93.5 % of crosslinks were considered satisfied in at least one state. **(D)** Clusters of overlength crosslinks from photo-crosslinking MS indicate conformational changes. RPN1 is particularly flexible in relation to the rest of the RP; in PDB IDs 7W3I and 7W37, different clusters of crosslink distance-restraints are satisfied. **(E)** A cluster of overlength crosslinks between RPN8 and RPN9 remains unsatisfied in all investigated proteasome structures. This suggests that an unsolved configuration exists between these subunits. **(F)** 276 distinct PPIs from RPN1 and RPN11 datasets combined at an estimated combined 7.7 % PPI-level FDR. Subunits of the 20S CP and the 19S RP are merged for simplicity. Each edge represents one or more crosslinks and line thickness scales with number of unique residue pairs per protein-protein interaction. The crosslink mapping to RAD23A or RAD23B is to a common peptide sequence prohibiting each to be distinguished. **(G)** 156 distinct protein-protein interactions discovered by crosslinking MS at 5 % FDR from the PSMB4 pulldown.

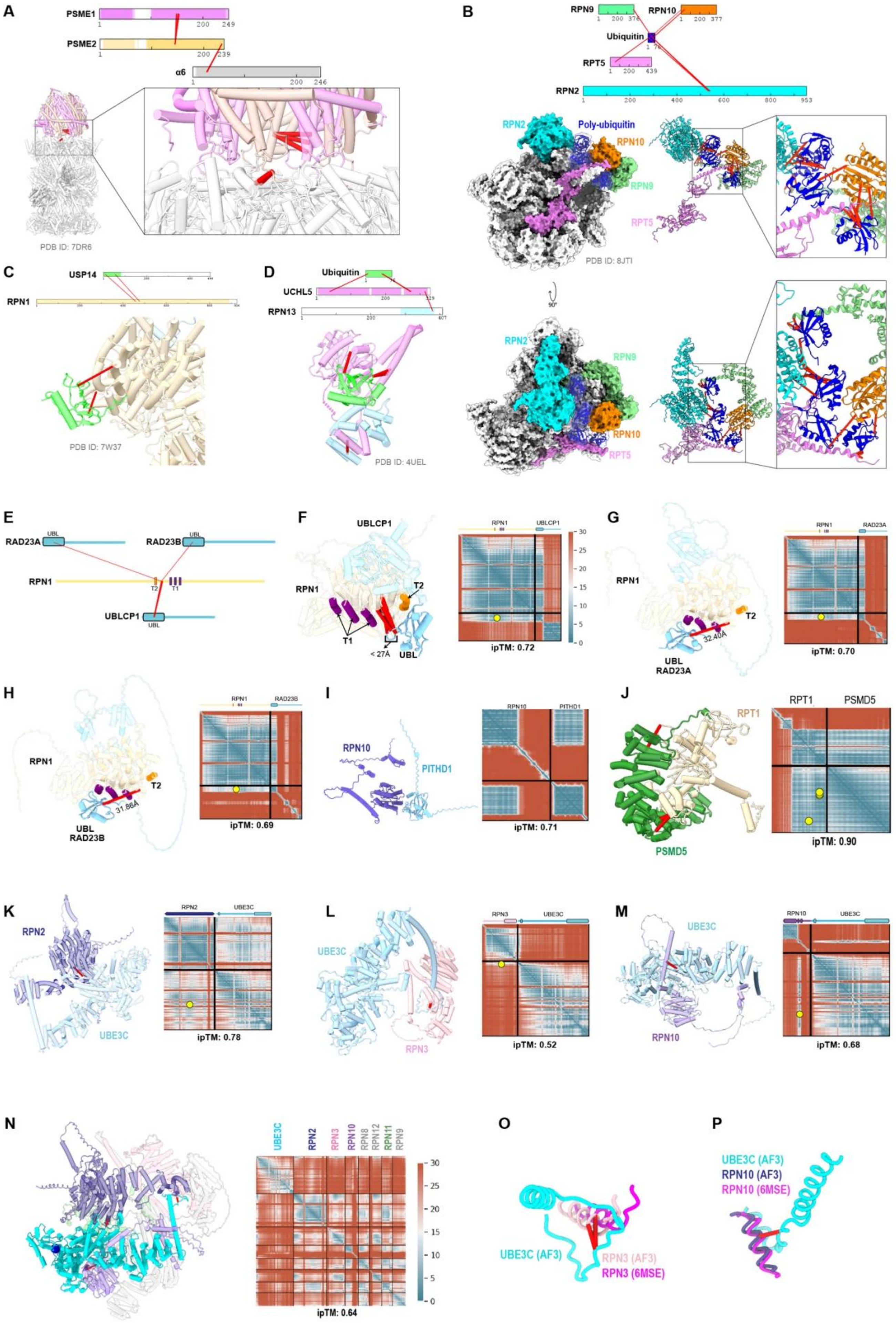
A
PSME1
PSME2
α6
PDB ID: 7DR6
B
RPN9
RPN10
Ubiquitin
RPT5
RPN2
Poly-ubiquitin
PDB ID: 8JTI
90°
C
USP14
RPN1
PDB ID: 7W37
D
Ubiquitin
UCHL5
RPN13
PDB ID: 4UEL
E
RAD23A
RAD23B
UBL
RPN1
T2
T1
UBLCP1
F
UBLCP1
RPN1
T1
T2
UBL
< 27Å
ipTM: 0.72
G
RPN1
RAD23A
T2
32.40Å
UBL
ipTM: 0.70
H
RPN1
RAD23B
T2
31.86Å
UBL
ipTM: 0.69
I
RPN10
PITHD1
ipTM: 0.71
J
RPT1
PSMD5
ipTM: 0.90
K
RPN2
UBE3C
ipTM: 0.78
L
UBE3C
RPN3
ipTM: 0.52
M
UBE3C
RPN10
ipTM: 0.68
N
UBE3C
RPN2
RPN3
RPN10
RPN8
RPN12
RPN11
RPN9
ipTM: 0.64
O
UBE3C (AF3)
RPN3 (AF3)
RPN3 (6MSE)
P
UBE3C (AF3)
RPN10 (AF3)
RPN10 (6MSE)

**Figure S4. Crosslinks on known and predicted structures for transiently associated proteins with proteasomes or their intrinsic subunits, related to Figure 2**. **(A)** Crosslinks on structure of the CP bound to the PSME1/2 alternate cap (PDB ID: 7DR6). **(B)** In situ-derived crosslinks on the structure of a K11/K48-branched ubiquitin chain on the 19S RP. Identified crosslinks with RPN2, RPN9, RPN10, and RPT5 agree with the available structure (PDB ID: 8JTI). **(C)** Crosslinks on the structure of USP14 bound to RPN1 (PDB ID: 7W37). **(D)** Crosslinks on the structure of the RPN13, UCHL5, and ubiquitin complex (PDB ID: 4UEL). **(E)** Bar representation of RPN1 and its crosslinked binders UBLCP1 and RAD23A/B with the crosslinked connections in red. The UBL domains and T1 and T2 sites are displayed and labeled. **(F-L)** AlphaFold-predicted as binary complexes (left) with ipTM scores >0.65, including **(F)** the UBLCP1 UBL domain bound to the RPN1 T2 site, RPN1 T1 bound to **(G)** RAD23A and **(H)** RAD23B UBL domains, **(I)** RPN10 VWA bound to PITHD1, **(J)** PSMD5 with RPT1, UBE3C bound to **(K)** RPN2, **(L)** RPN3, or **(M)** RPN10 VWA. No crosslinks were detected between RPN10 and PITHD1. (**N)** AlphaFold3 model comprising RPN2, RPN3, RPN8, RPN9, RPN10, RPN11, RPN12, and UBE3C that was used in Figure 2E. Crosslinked sites are depicted in red. Predicted alignment error (PAE) plots are included (right) showing residue-by-residue confidence scores of the predictions based on pairwise residue alignments with a yellow dot representing crosslinked residues. **(O-P)** Expanded regions of UBE3C (cyan) bound to RPN3 (aa 288-315, O) or RPN10 (aa 525-538, P) in the AlphaFold3 model of panel N (light pink in O and blue in P) superimposed as in Figure 2E onto a human 26S proteasome cryo-EM structure (PDB 6MSE, magenta in O and P). UBE3C residues 1 – 38 are displayed in (O) and 539 – 579 in (P).

A

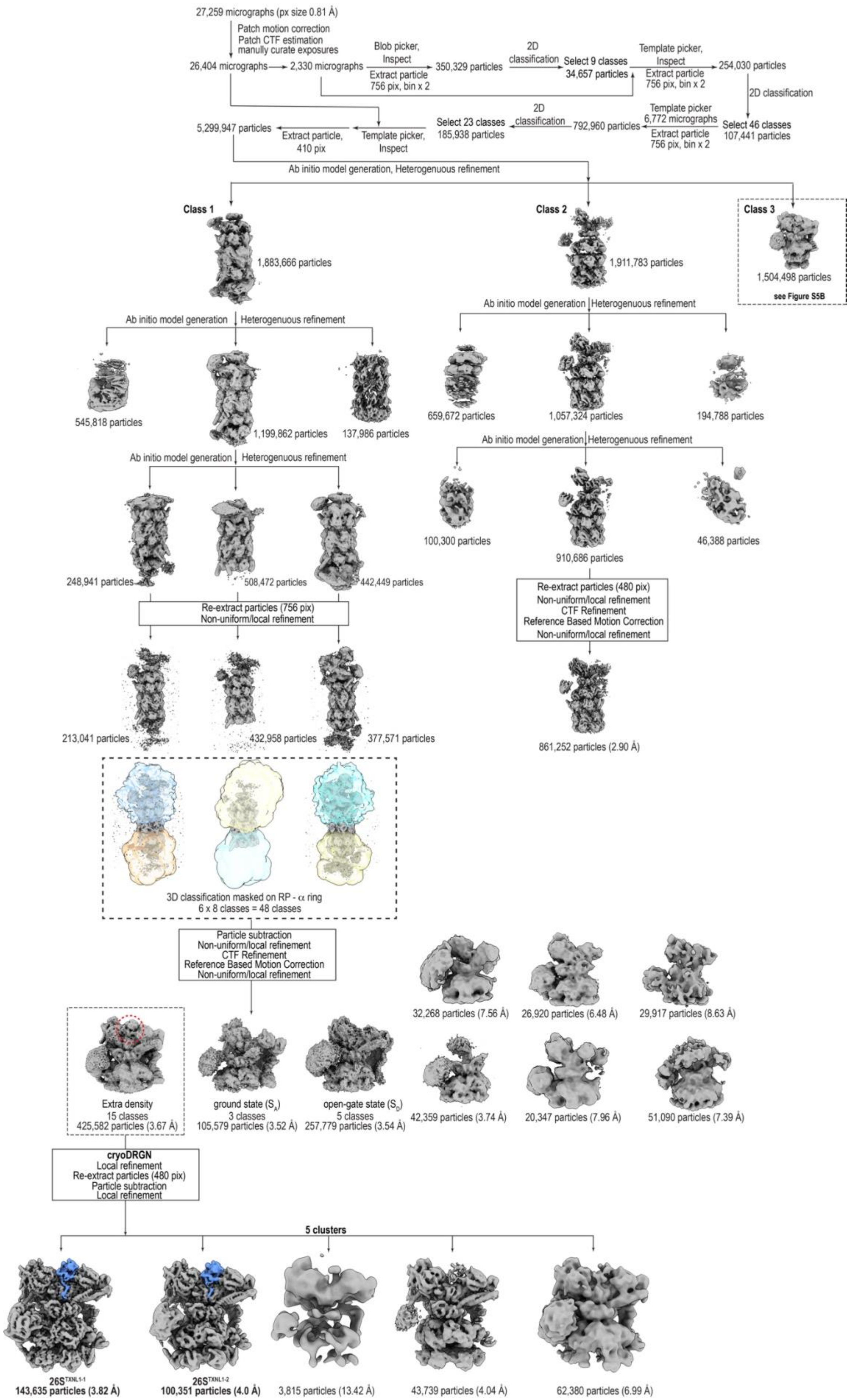

27,259 micrographs (px size 0.81 Å)
Patch motion correction
Patch CTF estimation
manully curate exposures
26,404 micrographs
2,330 micrographs
Blob picker, Inspect
Extract particle 756 pix, bin x 2
350,329 particles
2D classification
Select 9 classes
34,657 particles
Template picker, Inspect
Extract particle 756 pix, bin x 2
254,030 particles
2D classification
Select 46 classes
107,441 particles
Template picker
6,772 micrographs
Extract particle 756 pix, bin x 2
792,960 particles
2D classification
Select 23 classes
185,938 particles
Template picker, Inspect
Extract particle, 410 pix
5,299,947 particles
Ab initio model generation, Heterogenuous refinement
Class 1
1,883,666 particles
Class 2
1,911,783 particles
Class 3
1,504,498 particles
see Figure S5B
Ab initio model generation
Heterogenuous refinement
545,818 particles
1,199,862 particles
137,986 particles
Ab initio model generation
Heterogenuous refinement
248,941 particles
508,472 particles
442,449 particles
Re-extract particles (756 pix)
Non-uniform/local refinement
213,041 particles
432,958 particles
377,571 particles
3D classification masked on RP - α ring
6 x 8 classes = 48 classes
Particle subtraction
Non-uniform/local refinement
CTF Refinement
Reference Based Motion Correction
Non-uniform/local refinement
Extra density
15 classes
425,582 particles (3.67 Å)
ground state (S_A)
3 classes
105,579 particles (3.52 Å)
open-gate state (S_D)
5 classes
257,779 particles (3.54 Å)
32,268 particles (7.56 Å)
26,920 particles (6.48 Å)
29,917 particles (8.63 Å)
42,359 particles (3.74 Å)
20,347 particles (7.96 Å)
51,090 particles (7.39 Å)
Ab initio model generation
Heterogenuous refinement
659,672 particles
1,057,324 particles
194,788 particles
Ab initio model generation
Heterogenuous refinement
100,300 particles
910,686 particles
46,388 particles
Re-extract particles (480 pix)
Non-uniform/local refinement
CTF Refinement
Reference Based Motion Correction
Non-uniform/local refinement
861,252 particles (2.90 Å)
cryoDRGN
Local refinement
Re-extract particles (480 pix)
Particle subtraction
Local refinement
5 clusters
26S^TXNL1-1
143,635 particles (3.82 Å)
26S^TXNL1-2
100,351 particles (4.0 Å)
3,815 particles (13.42 Å)
43,739 particles (4.04 Å)
62,380 particles (6.99 Å)

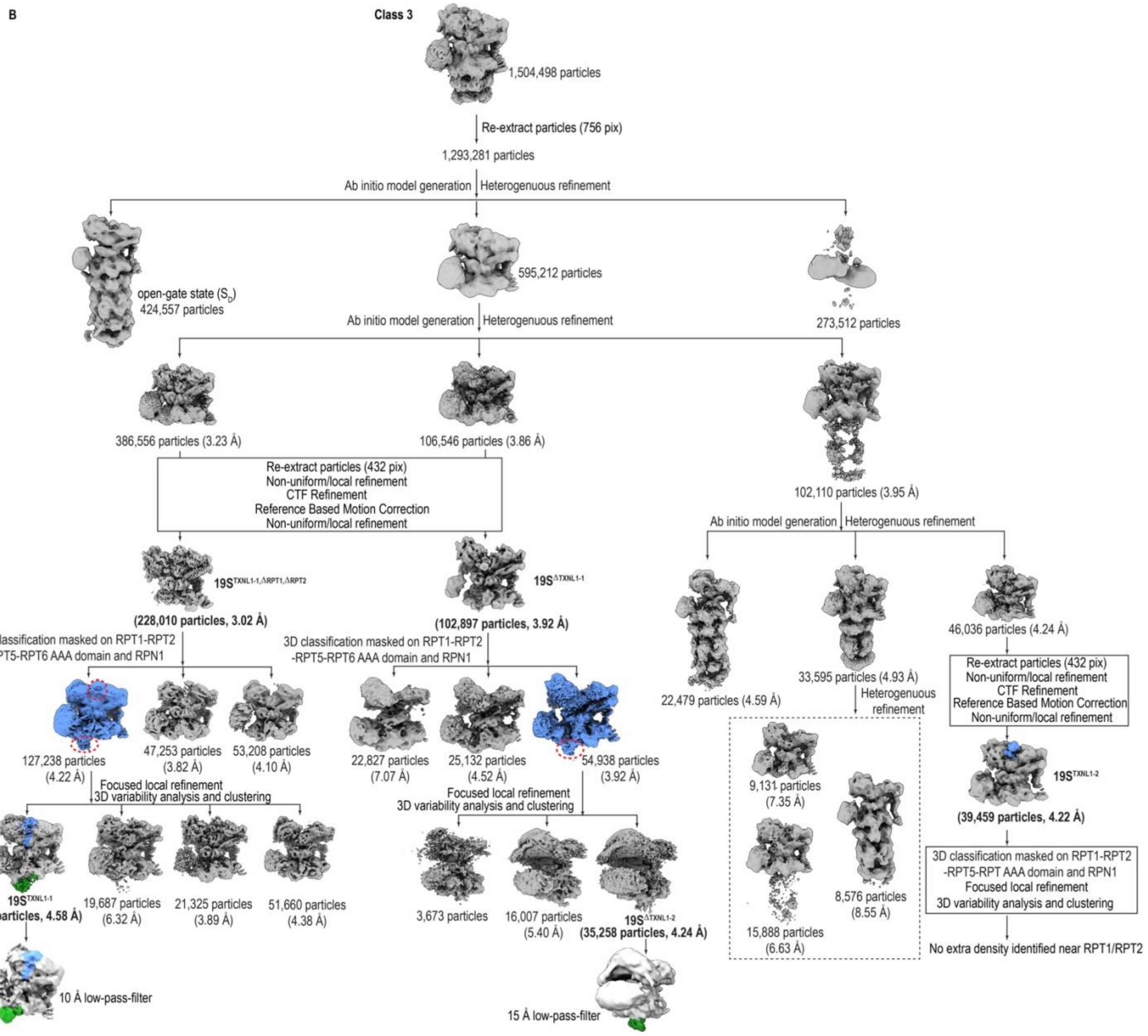

B
Class 3
1,504,498 particles
Re-extract particles (756 pix)
1,293,281 particles
Ab initio model generation
Heterogenuous refinement
open-gate state ($S_D$)
424,557 particles
595,212 particles
273,512 particles
Ab initio model generation
Heterogenuous refinement
386,556 particles (3.23 Å)
106,546 particles (3.86 Å)
102,110 particles (3.95 Å)
Re-extract particles (432 pix)
Non-uniform/local refinement
CTF Refinement
Reference Based Motion Correction
Non-uniform/local refinement
$19S^{TXNL1-1,\Delta RPT1,\Delta RPT2}$
(228,010 particles, 3.02 Å)
$19S^{\Delta TXNL1-1}$
(102,897 particles, 3.92 Å)
Ab initio model generation
Heterogenuous refinement
22,479 particles (4.59 Å)
33,595 particles (4.93 Å)
46,036 particles (4.24 Å)
3D classification masked on RPT1-RPT2
-RPT5-RPT6 AAA domain and RPN1
3D classification masked on RPT1-RPT2
-RPT5-RPT6 AAA domain and RPN1
127,238 particles
(4.22 Å)
47,253 particles
(3.82 Å)
53,208 particles
(4.10 Å)
22,827 particles
(7.07 Å)
25,132 particles
(4.52 Å)
54,938 particles
(3.92 Å)
Heterogenuous
refinement
9,131 particles
(7.35 Å)
15,888 particles
(6.63 Å)
8,576 particles
(8.55 Å)
Re-extract particles (432 pix)
Non-uniform/local refinement
CTF Refinement
Reference Based Motion Correction
Non-uniform/local refinement
$19S^{TXNL1-2}$
(39,459 particles, 4.22 Å)
Focused local refinement
3D variability analysis and clustering
Focused local refinement
3D variability analysis and clustering
$19S^{TXNL1-1}$
(34,566 particles, 4.58 Å)
19,687 particles
(6.32 Å)
21,325 particles
(3.89 Å)
51,660 particles
(4.38 Å)
3,673 particles
16,007 particles
(5.40 Å)
$19S^{\Delta TXNL1-2}$
(35,258 particles, 4.24 Å)
3D classification masked on RPT1-RPT2
-RPT5-RPT AAA domain and RPN1
Focused local refinement
3D variability analysis and clustering
No extra density identified near RPT1/RPT2
10 Å low-pass-filter
15 Å low-pass-filter

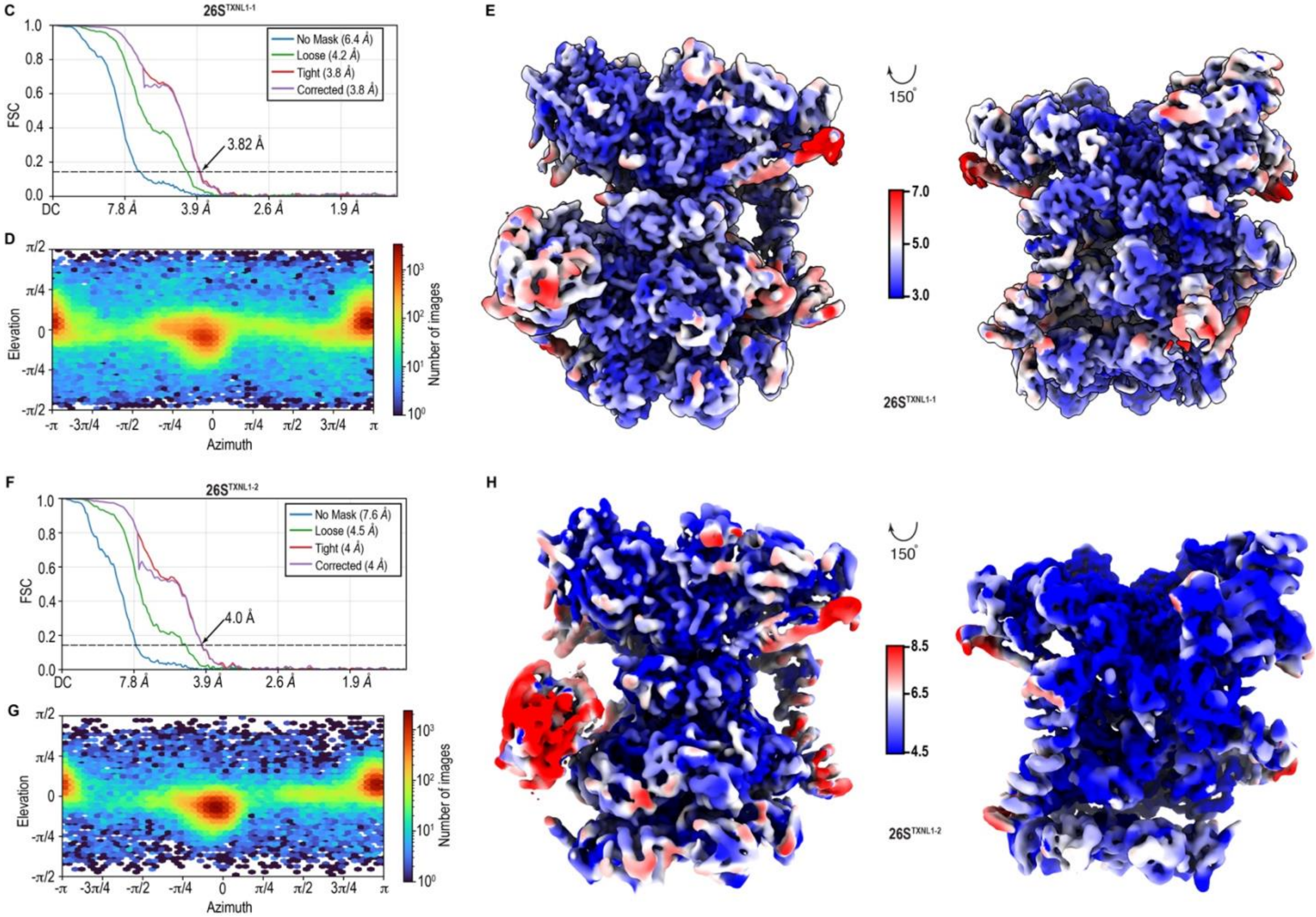
C
26S^TXNL1-1
No Mask (6.4 Å)
Loose (4.2 Å)
Tight (3.8 Å)
Corrected (3.8 Å)
3.82 Å
FSC
DC 7.8 Å 3.9 Å 2.6 Å 1.9 Å
D
Elevation
Azimuth
Number of images
E
150°
7.0
5.0
3.0
26S^TXNL1-1
F
26S^TXNL1-2
No Mask (7.6 Å)
Loose (4.5 Å)
Tight (4 Å)
Corrected (4 Å)
4.0 Å
FSC
DC 7.8 Å 3.9 Å 2.6 Å 1.9 Å
G
Elevation
Azimuth
Number of images
H
150°
8.5
6.5
4.5
26S^TXNL1-2

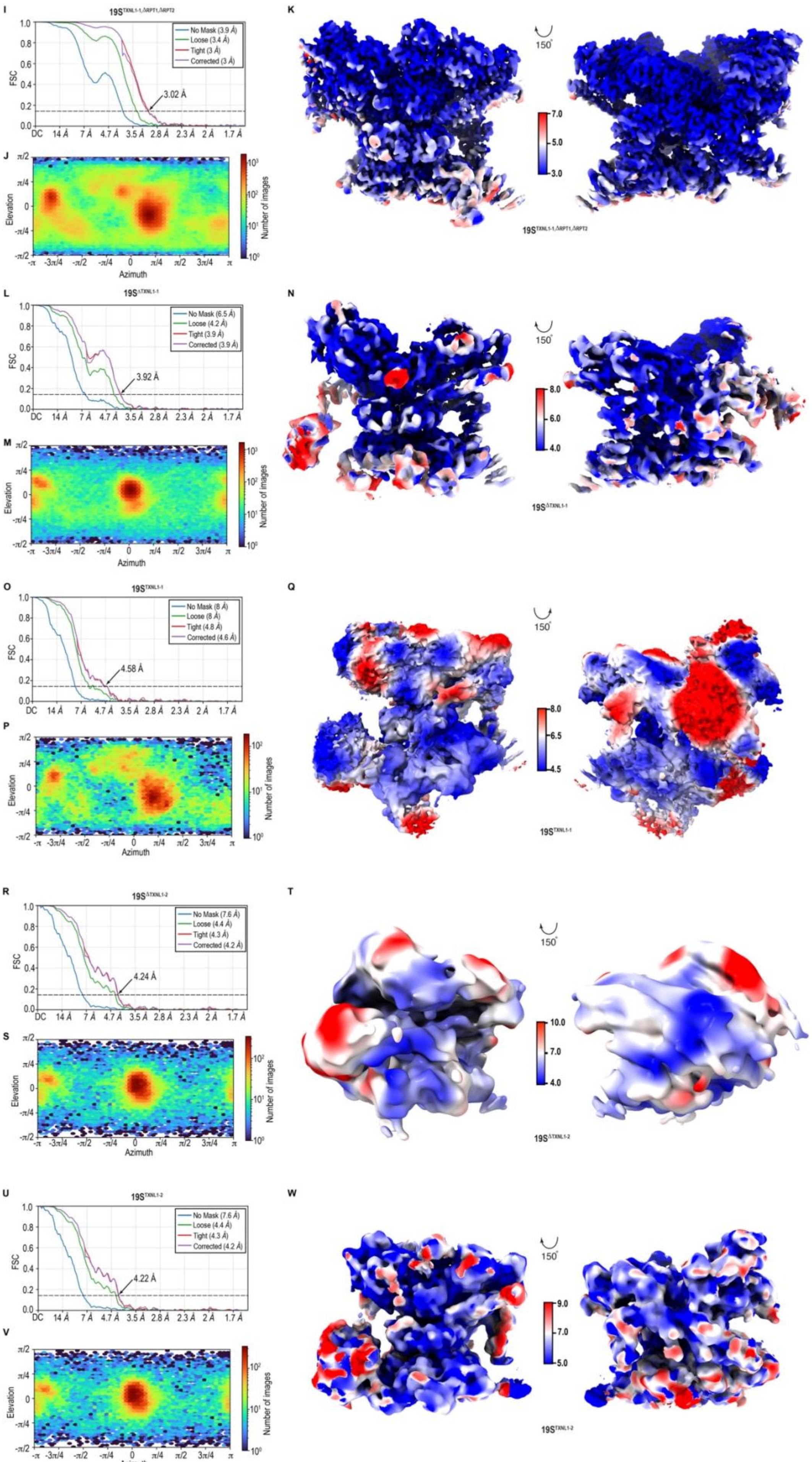

I
$19S^{TXNL1\text{-}1,\Delta RPT1,\Delta RPT2}$
No Mask (3.9 Å)
Loose (3.4 Å)
Tight (3 Å)
Corrected (3 Å)
3.02 Å
FSC
J
Elevation
Azimuth
Number of images
K
150°
$19S^{TXNL1\text{-}1,\Delta RPT1,\Delta RPT2}$
L
$19S^{\Delta TXNL1\text{-}1}$
No Mask (6.5 Å)
Loose (4.2 Å)
Tight (3.9 Å)
Corrected (3.9 Å)
3.92 Å
M
N
$19S^{\Delta TXNL1\text{-}1}$
O
$19S^{TXNL1\text{-}1}$
No Mask (8 Å)
Loose (8 Å)
Tight (4.8 Å)
Corrected (4.6 Å)
4.58 Å
P
Q
$19S^{TXNL1\text{-}1}$
R
$19S^{\Delta TXNL1\text{-}2}$
No Mask (7.6 Å)
Loose (4.4 Å)
Tight (4.3 Å)
Corrected (4.2 Å)
4.24 Å
S
T
$19S^{\Delta TXNL1\text{-}2}$
U
$19S^{TXNL1\text{-}2}$
No Mask (7.6 Å)
Loose (4.4 Å)
Tight (4.3 Å)
Corrected (4.2 Å)
4.22 Å
V
W
$19S^{TXNL1\text{-}2}$

**Figure S5. Cryo-EM workflow for proteasome complexes purified using an RPN1-affinity tag, related to Figure 3, 4, and 6 (A)** Processing scheme for the cryo-EM classification process for the full dataset showing reconstructions following refinement of $26S^{TXNL1-1}$ and $26S^{TXNL1-2}$. Data were collected on proteasomes isolated from RPN1-tagged HCT116 cells by using a Talos Arctica G2 electron microscope (Thermo Fisher Scientific, Inc) equipped with a K3 direct electron detector (Gatan) and an energy filter, operating at 200 kV in super resolution mode (pixel size 0.405 Å/pixel, x 100,000 nominal magnification). **(B)** Processing scheme for the classification of class 3 particles from panel A showing reconstructions following refinement of $19S^{TXNL1-1}$, $19S^{TXNL1-2}$, $19S^{TXNL1-1,\Delta RPT1, \Delta RPT2}$, $19S^{\Delta TXNL1-1}$, and $19S^{\Delta TXNL1-2}$. **(C-W)** GSFSC plots (C, F, I, L, O, R, and U), viewing direction distribution plots (D, G, J, M, P, S, and V), and cryo-EM density maps displaying local resolution as calculated by cryoSPARC or Phenix suite (E, H, K, N, Q, T, and W) for the refined maps of $26S^{TXNL1-1}$ (C, D, and E), $26S^{TXNL1-2}$ (F, G, and H), $19S^{TXNL1-1,\Delta RPT1, \Delta RPT2}$ (I, J, and K), $19S^{\Delta TXNL1-1}$ (L, M, and N), $19S^{TXNL1-1}$ (O, P, and Q), $19S^{\Delta TXNL1-2}$ (R, S, and T), and $19S^{TXNL1-2}$ (U, V, and W). FSC value of 0.143 is indicated by a black dashed line in C, F, I, L, O, R, and U.

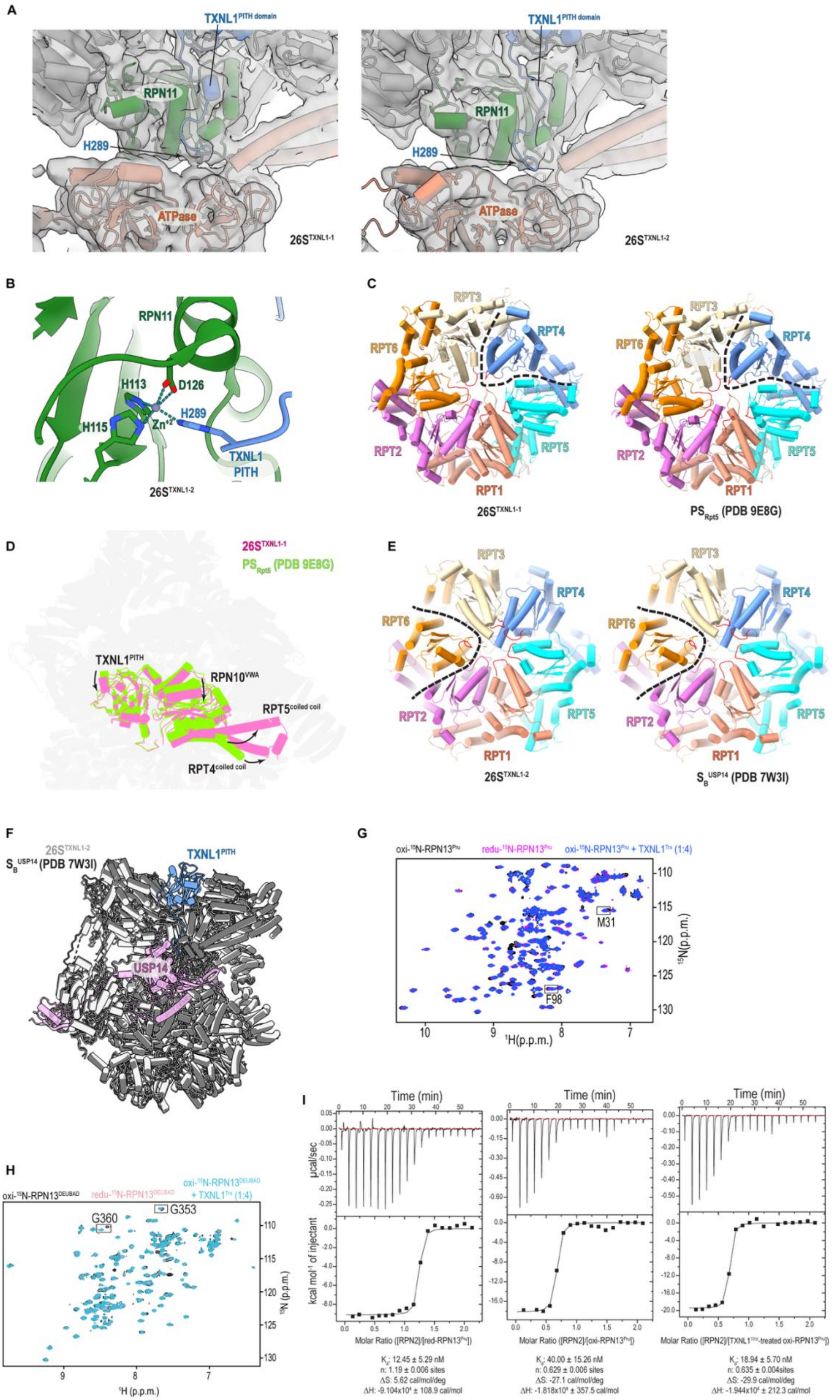

A
TXNL1PITH domain
RPN11
H289
ATPase
26STXNL1-1
TXNL1PITH domain
RPN11
H289
ATPase
26STXNL1-2
B
RPN11
H113
D126
H115
Zn+2
H289
TXNL1 PITH
26STXNL1-2
C
RPT3
RPT4
RPT6
RPT2
RPT5
RPT1
26STXNL1-1
PSRpt5 (PDB 9E8G)
D
26STXNL1-1
PSRpt5 (PDB 9E8G)
TXNL1PITH
RPN10VWA
RPT5coiled coil
RPT4coiled coil
E
RPT3
RPT4
RPT6
RPT2
RPT5
RPT1
26STXNL1-2
SBUSP14 (PDB 7W3I)
F
26STXNL1-2
SBUSP14 (PDB 7W3I)
TXNL1PITH
USP14
G
oxi-15N-RPN13Pru
redu-15N-RPN13Pru
oxi-15N-RPN13Pru + TXNL1Trx (1:4)
M31
F98
15N(p.p.m.)
1H(p.p.m.)
H
oxi-15N-RPN13DEUBAD
redu-15N-RPN13DEUBAD
oxi-15N-RPN13DEUBAD + TXNL1Trx (1:4)
G360
G353
15N (p.p.m.)
1H (p.p.m.)
I
Time (min)
µcal/sec
kcal mol-1 of injectant
Molar Ratio ([RPN2]/[red-RPN13Pru])
Molar Ratio ([RPN2]/[oxi-RPN13Pru])
Molar Ratio ([RPN2]/[TXNL1TRX-treated oxi-RPN13Pru])
Kd: 12.45 ± 5.29 nM
n: 1.19 ± 0.006 sites
ΔS: 5.62 cal/mol/deg
ΔH: -9.104x10⁴ ± 108.9 cal/mol
Kd: 40.00 ± 15.26 nM
n: 0.629 ± 0.006 sites
ΔS: -27.1 cal/mol/deg
ΔH: -1.818x10⁴ ± 357.5 cal/mol
Kd: 18.94 ± 5.70 nM
n: 0.635 ± 0.004sites
ΔS: -29.9 cal/mol/deg
ΔH: -1.944x10⁴ ± 212.3 cal/mol

**Figure S6. TXNL1 activities at the proteasome, related to Figure 3 and 4**. **(A)** Density maps (grey) for 26S$^{TXNL1-1}$ (left) and 26S$^{TXNL1-2}$ (right) focused on the C-terminus with fitted cartoon models. The coloring scheme follows Figure 3A. **(B)** Expanded view of 26S$^{TXNL1-2}$ to display TXNL1 PITH domain and C-terminal residue H289 coordination with the RPN11 $Zn^{+2}$ (purple sphere) and its additional coordination with H113, H115 and D126 of RPN11. **(C and E)** Top views of the ATPase large AAA+ subdomains in aligned 26S$^{TXNL1-1}$ (C, left) and $PS_{Rpt5}$ (PDB 9E8G, B, right), or 26S$^{TXNL1-2}$ (E, left) and $S_B^{USP14}$ (PDB 7WSI, E, right), with the same color scheme as used in Figure 3E and 3F. The reduced interleaving between RPT4 (C) or RPT6 (E) and the neighboring RPT subunits is highlighted as black dashed lines. **(D)** Superimposed structures of 26S$^{TXNL1-1}$ and $PS_{Rpt5}$ (PDB 9E8G), with the TXNL1 PITH domain, RPN10 VWA domain, and RPT4:RPT5 coiled coil highlighted in pink (26S$^{TXNL1-1}$) and green ($PS_{Rpt5}$). The shift or rotation of these regions in 26S$^{TXNL1-1}$ against $PS_{Rpt5}$ is indicated by black arrows. **(F)** Superimposed structures of 26S$^{TXNL1-2}$ and USP14-bound 26S proteasome $S_B^{USP14}$ (PDB 7WSI) with TXNL1$^{PITH}$ and USP14 colored in blue and pink, respectively. The other subunits are colored in light grey (26S$^{TXNL1-2}$) and dark grey ($S_B^{USP14}$). **(G)** Overlaid $^{1}$H-$^{15}$N HSQC spectra of reduced $^{15}$N-RPN13$^{Pru}$ (22 μM, redu, pink), oxidized $^{15}$N-RPN13$^{Pru}$ (22 μM, oxi, black) and oxidized $^{15}$N-RPN13$^{Pru}$ with 4-fold molar excess of TXNL1$^{Trx}$ (88 μM, blue). Enlarged regions for residues M31 and F98, highlighted in black boxes, are shown in Figure 4C. **(H)** Overlaid $^{1}$H-$^{15}$N HSQC spectra of reduced (redu) $^{15}$N-RPN13$^{DEUBAD}$ (25 μM, rose-pink), oxidized (oxi) $^{15}$N-RPN13$^{DEUBAD}$ (25 μM, black) and oxidized RPN13$^{DEUBAD}$ with 4-fold molar excess of TXNL1$^{Trx}$ (100 μM, cyan blue). Enlarged region for residues G353 and G360, highlighted in black boxes, are shown in Figure 4D. All spectra were acquired at 600 MHz and 10 °C with NMR buffer containing 20 mM sodium phosphate and 50 mM NaCl (pH 6.0). In addition, 1 mM DTT was added for samples under reducing condition. **(I)** Left panel: 156 µM RPN2 peptide (940-953) in reducing ITC buffer (20 mM sodium phosphate, 50 mM NaCl, 10 mM βME, pH 6.0) was injected into a calorimeter cell containing reduced 15.6 µM RPN13$^{Pru}$. Middle panel: 205.7 μM RPN2 peptide in non-reducing ITC buffer (20 mM sodium phosphate, 50 mM NaCl, pH 6.0) was injected into a calorimeter cell containing oxidized 20.57 µM RPN13$^{Pru}$. Right panel: 160 μM RPN2 peptide in non-reducing ITC buffer was injected into a calorimeter cell containing 16 μM oxidized RPN13$^{Pru}$ pre-incubated with 3-fold molar excess of TXNL1$^{Trx}$ and separated by size exclusion chromatography. All binding isotherms (top) were integrated to yield the change in enthalpy as a function of RPN2 peptide addition. The data were fit to a “One Set of Sites” binding model with the indicated thermodynamic values (bottom). All ITC experiment were performed at 25 °C.

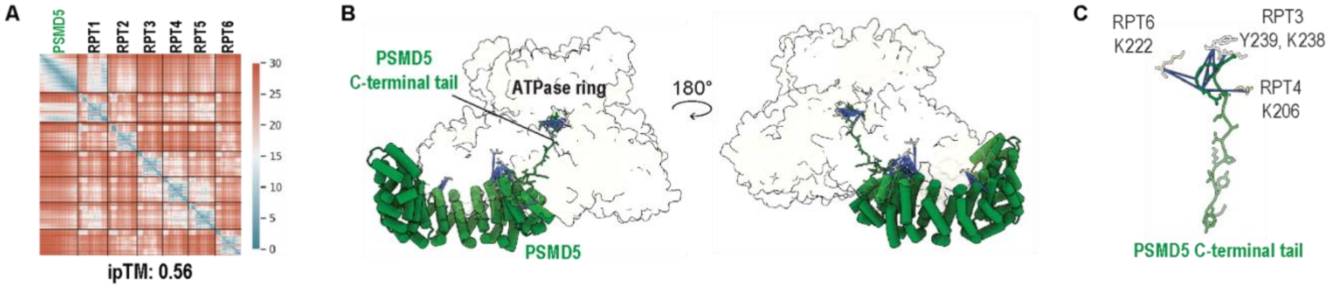

**Figure S7. AlphaFold predicts a confidently scored model of PSMD5 bound to the proteasome ATPase ring, related to Figure 5. (A)** Predicted Alignment Error (PAE) plot of a selected AlphaFold prediction. **(B)** AlphaFold3 predicted model of PSMD5 (green, cartoon view) bound to the ATPase ring (white, surface rendering) with crosslinks displayed by blue lines. **(C)** Expanded view an AlphaFold3 model centered on the PSMD5 C-terminal tail (green) to display its crosslinks (blue lines) with RPT6, RPT3, and RPT4. The sidechain of residues K222 (RPT6), K238 (RPT3), Y239 (RPT3), and G206 (RPT4) are displayed.

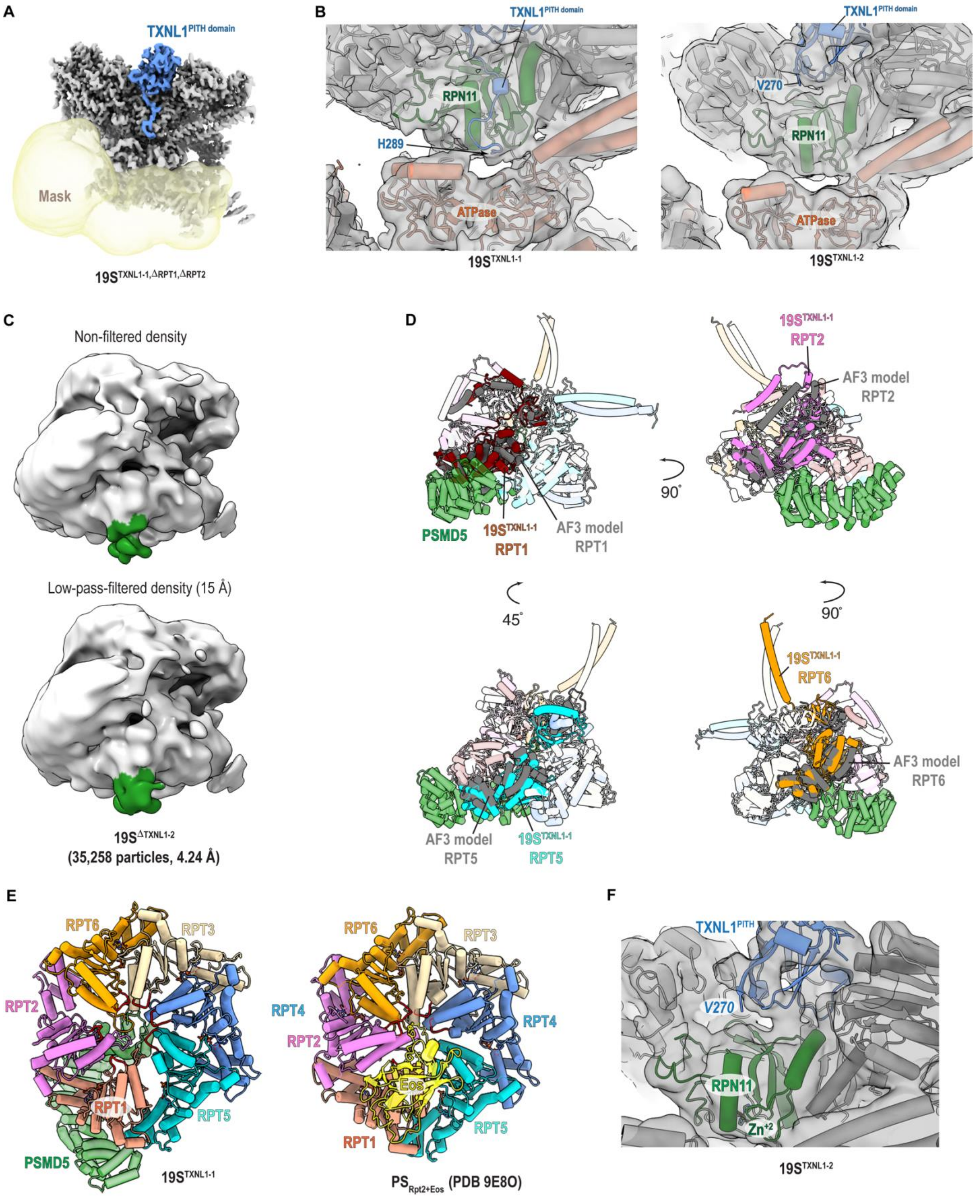
A
TXNL1 PITH domain
Mask
19S TXNL1-1,ΔRPT1,ΔRPT2
B
TXNL1 PITH domain
RPN11
H289
ATPase
19S TXNL1-1
TXNL1 PITH domain
V270
RPN11
ATPase
19S TXNL1-2
C
Non-filtered density
Low-pass-filtered density (15 Å)
19S ΔTXNL1-2
(35,258 particles, 4.24 Å)
D
PSMD5
19S TXNL1-1
RPT1
AF3 model
RPT1
90°
19S TXNL1-1
RPT2
AF3 model
RPT2
45°
90°
AF3 model
RPT5
19S TXNL1-1
RPT5
19S TXNL1-1
RPT6
AF3 model
RPT6
E
RPT6
RPT3
RPT2
RPT4
RPT1
RPT5
PSMD5
19S TXNL1-1
RPT6
RPT3
RPT4
RPT2
RPT4
Eos
RPT1
RPT5
PS Rpt2+Eos (PDB 9E8O)
F
TXNL1 PITH
V270
RPN11
Zn+2
19S TXNL1-2

**Figure S8. 19S RP with TXNL1 and PSMD5 bound, related to Figure 6**. **(A)** A mask (yellow map) focused on the RPT1-RPT2-RPT5-RPT6 AAA domain and RPN1 was created and used for 3D classification, 3D variability analyses and local refinement of $19S^{TXNL1\text{-}1,\Delta RPT1,\ \Delta RPT2}$. **(B**) Density maps (grey) for $19S^{TXNL1\text{-}1}$ (left) and $19S^{TXNL1\text{-}2}$ (right) focused on the C-terminus with fitted cartoon models. The coloring scheme follows Figure 6A. **(C)** Extra density (green) is detected in the non-filtered (top) and low-pass-filtered (15 Å, bottom) $19S^{\Delta TXNL1\text{-}2}$ map near the ATPase ring. **(D)** Expanded views of the ATPase ring and PSMD5 in the aligned $19S^{TXNL1\text{-}1}$ and AlphaFold3 model structures. Alignment was based on PSMD5. $19S^{TXNL1\text{-}1}$ uses the same color scheme as in Figure 6E. RPT subunits in the AlphaFold3 model are colored in white, except for the one that is used for comparison with $19S^{TXNL1\text{-}1}$, which is highlighted dark grey. **(E)** Top views of the ATPase large AAA+ subdomains in aligned $19S^{TXNL1\text{-}1}$ (left) and $PS_{Rpt2+Eos}$ (right) structures, with the same color scheme as used in Figure 6G. PSMD5 and Eos are colored in green and yellow, respectively. **(F)** Expanded view of the $19S^{TXNL1\text{-}2}$ structure to demonstrate that the 20 amino acids after V270 in the TXNL1 C-terminal tail (blue) is not detected and not interacting with the RPN11 $Zn^{+2}$ (green). The density map of $19S^{TXNL1\text{-}2}$ is displayed in grey.

**Table S1. Guide RNAs tested in 293T cells**

| Name | Gene | Human/ Mouse | Description | Target |
|---|---|---|---|---|
| IVT-2994 | PSMD14 | Human | PSMD14-Cterm_FASTA:75-97 | TTAGGACCCCAAACGTCATTTGG |
| IVT-2995* | PSMD14 | Human | PSMD14-Cterm_FASTA:81-103 | GTTCCTCCAAATGACGTTTGGGG |
| IVT-2996 | PSMD14 | Human | PSMD14-Cterm_FASTA:83-105 | ATGTTCCTCCAAATGACGTTTGG |
| IVT-2997* | PSMD14 | Human | PSMD14-Cterm_FASTA:248-270 | GGGACCTCTGAAGGTGTACTTG G |
| JT-IVT-44* | PSMB4 | Human | PSMB4_Cterm_83forw | TGGACAGTACAGCTATTTTTAGG |
| JT-IVT-45 | PSMB4 | Human | PSMB4_Cterm_127rev | TCATTCAAAGCCACTAAAAAAGG |
| JT-IVT-46 | PSMB4 | Human | PSMB4_Cterm_166rev | AAGTAGGGCAACTTCAGTTCTGG |
| JT-IVT-47 | PSMB4 | Human | PSMB4_Cterm_202forw | CCTACTTTTAACTTTGAACTTGG |
| JT-IVT-48* | PSMB4 | Human | PSMB4_Cterm_181rev | CAAGTTCAAAGTTAAAAGTAGGG |
| JT-IVT-49 | PSMB4 | Human | PSMB4_Cterm_182rev | CCAAGTTCAAAGTTAAAAGTAGG |

* Guide RNAs used for knock-in experiments.

**Table S2. Chromosomal locations of PSMD14 and PSMB4 as per GRCh38/Hg38**

| Name | Regions | Chromosomal location |
|---|---|---|
| hRpn11-01 | IVT-2995 | chr2:161,411,304-161,411,326 |
| hRpn11-02 | IVT-2997 | chr2:161,411,471-161,411,493 |
| PSMB4-01 | JT-IVT-44 | chr1:151,401,743-151,401,765 |
| PSMB4-02 | JT-IVT-48 | chr1:151,401,861-151,401,883 |

**Table S3. Oligos for guide RNAs**

| Name | Forward oligo | Reverse oligo |
|---|---|---|
| IVT-2995 | CACCGGTTCCTCCAAATGACGTTTG | AAACCAAACGTCATTTGGAGGAACC |
| IVT-2997 | ACCGGGGACCTCTGAAGGTGTACT GT | TAAAACAGTACACCTTCAGAGGTCCC |
| JT-IVT-44 | CACCGTGGACAGTACAGCTATTTTT | AAACAAAAATAGCTGTACTGTCCAC |
| JT-IVT-48 | ACCGCAAGTTCAAAGTTAAAAGTA GT | TAAAAC ACTTTTAACTTTGAACTTG |

### Table S4. Details of the primers used in the study

| Primer Name | Primer sequence |
|---|---|
| PSMD14-GT-V1-F | GTTAGAAAAGCATGGCCAAG |
| PSMD14-GT-V1-R | CTATTAGAAAACTGGGTAGTC |
| PSMD14-GT-V2-F | GTTAGGAGTCTTGGGTATTT |
| PSMD14-GT-V2-R | AGATGAGACCAAACAATAGC |
| PSMB4-GT-V1-F | CGCTATTACTGGGCAGATGG |
| PSMB4-GT-V1-R | CTGCCTGCATGGCTTAGTAG |
| PSMB4-GT-V2-F | GTGGTTGTCTGCTTTTCTCC |
| PSMB4-GT-V2-R | CTGTGTTGAGACTGAATGTG |

### Table S5. Statistics for cryo-EM analyses of 26S$^{TXNL1-1}$, 26S$^{TXNL1-2}$, 19S$^{TXNL1-1}$, 19S$^{TXNL1-2}$, and 19S$^{TXNL1-1,\Delta RPT1\ \Delta RPT2}$.

| | 26S$^{TXNL1-1}$ | 26S$^{TXNL1-2}$ | 19S$^{TXNL1-1}$ | 19S$^{TXNL1-2}$ | 19S$^{TXNL1-1,\Delta RPT1\ \Delta RPT2}$ |
|---|---|---|---|---|---|
| **Data collection and image processing** | | | | | |
| Microscope | Talos Arctica G2 | | | | |
| Camera | K3 | | | | |
| Magnification | 100,000 | | | | |
| Voltage (kV) | 200 | | | | |
| Electron exposure ($e^-$/Å$^2$) | 55.6 | | | | |
| Defocus range (μm) | 0.8 - 2.0 | | | | |
| Pixel size (Å) | 0.81 | | | | |
| Symmetry imposed | C1 | | | | |
| Total micrographs | 27,259 | | | | |
| Initial particle images | 5,299,947 | | | | |
| Final particle images | 143,635 | 100,351 | 34,566 | 39,459 | 228,010 |
| Map resolution (Å) | 3.82 | 4.0 | 4.58 | 4.22 | 3.02 |
| FSC threshold | 0.143 | 0.143 | 0.143 | 0.143 | 0.143 |
| Map sharpening *B* factor (Å$^2$) | -58.0 | -41.8 | -147.8 | -47.3 | -88.9 |
| EMDB code | 71740 | 71741 | 71810 | 71737 | 71813 |
| **Model building and refinement** | | | | | |
| Initial model used (PDB ID) | 7W3I,1WWY,1GH2 | 7W3I, 1WWY | 7W3I, 1WWY | 7W3I, 1WWY | 7W3I |
| Model resolution (Å) | 4.14 | 4.40 | 4.87 | 4.67 | 3.99 |
| FSC threshold | 0.143 | 0.143 | 0.143 | 0.143 | 0.143 |
| *Model composition* | | | | | |
| Non-hydrogen atoms | 71045 | 69593 | 59237 | 53748 | 41598 |
| Amino acid residues | 8972 | 8790 | 7457 | 6756 | 5206 |
| Protein molecules | 26 | 26 | 20 | 19 | 16 |
| *Real-space correlation* | | | | | |
| CC (volume) | 0.73 | 0.70 | 0.58 | 0.64 | 0.56 |
| CC (mask) | 0.75 | 0.73 | 0.61 | 0.66 | 0.57 |

| | | | | | |
|---|---|---|---|---|---|
| *RMS deviations* | | | | | |
| Bond lengths (Å) | 0.007 | 0.007 | 0.006 | 0.008 | 0.006 |
| Bond angles (°) | 1.384 | 1.468 | 1.311 | 1.442 | 1.310 |
| *Validation* | | | | | |
| MolProbity score | 1.75 | 2.12 | 1.67 | 2.13 | 1.64 |
| Clash score | 6.05 | 7.82 | 5.16 | 11.12 | 5.45 |
| Rotamers outliers (%) | 0.30 | 1.86 | 0.29 | 0.59 | 0.26 |
| CaBLAM outliers (%) | 4.23 | 5.27 | 3.16 | 6.04 | 2.89 |
| $C\beta$ outliers (%) | 0.02 | 0.01 | 0.00 | 0.00 | 0.00 |
| *Ramachandran plot* | | | | | |
| Favored (%) | 93.60 | 92.07 | 94.22 | 89.15 | 94.94 |
| Allowed (%) | 6.40 | 7.93 | 5.78 | 10.85 | 5.06 |
| Outliers (%) | 0.00 | 0.00 | 0.00 | 0.00 | 0.00 |
| PDB code | 9PMO | 9PMQ | 9PRO | 9PMJ | 9PRT |

**Table S6. Statistics for cryo-EM analyses of $19S^{\Delta TXNL1\text{-}1}$ and $19S^{\Delta TXNL1\text{-}2}$.**

| | **$19S^{\Delta TXNL1\text{-}1}$** | **$19S^{\Delta TXNL1\text{-}2}$** |
|---|---|---|
| **Data collection and image processing** | | |
| Microscope | Talos Arctica G2 | |
| Camera | K3 | |
| Magnification | 100,000 | |
| Voltage (kV) | 200 | |
| Electron exposure ($e^-$/Å$^2$) | 55.6 | |
| Defocus range (µm) | 0.8 - 2.0 | |
| Pixel size (Å) | 0.81 | |
| Symmetry imposed | C1 | |
| Total micrographs | 27,259 | |
| Initial particle images | 5,299,947 | |
| Final particle images | 102,897 | 35,258 |
| Map resolution (Å) | 3.92 | 4.24 |
| FSC threshold | 0.143 | 0.143 |
| Map sharpening *B* factor (Å$^2$) | -57.5 | -85.2 |
| EMDB code | 71791 | 71795 |
